# Human iPSC-derived neural stem cells display a radial glia-like signature *in vitro* and favorable long-term safety in transplanted mice

**DOI:** 10.1101/2023.08.04.551937

**Authors:** Marco Luciani, Chiara Garsia, Stefano Beretta, Luca Petiti, Clelia Peano, Ivan Merelli, Ingrid Cifola, Annarita Miccio, Vasco Meneghini, Angela Gritti

## Abstract

Human induced pluripotent stem cell-derived neural stem/progenitor cells (hiPSC-NSCs) are a promising source for cell therapy approaches to treat neurodegenerative and demyelinating disorders. Despite ongoing efforts to characterize hiPSC-derived cells *in vitro* and *in vivo*, we lack comprehensive genome- and transcriptome-wide studies addressing hiPSC-NSC identity and safety, which are critical for establishing accepted criteria for prospective clinical applications.

Here, we evaluated the transcriptional and epigenetic signatures of hiPSCs and differentiated hiPSC-NSC progeny, finding that the hiPSC-to-NSC transition results in a complete loss of pluripotency and the acquisition of a radial glia-associated transcriptional signature. Importantly, hiPSC-NSCs share with somatic human fetal NSCs (hfNSCs) the main transcriptional and epigenetic patterns associated with NSC-specific biology. *In vivo*, long-term observation (up to 10 months) of mice intracerebrally transplanted as neonates with hiPSC-NSCs showed robust engraftment and widespread distribution of human cells in the host brain parenchyma. Engrafted hiPSC-NSCs displayed multilineage potential and preferentially generated glial cells. No hyperproliferation, tumor formation, or expression of pluripotency markers was observed. Finally, we identified a novel role of the Sterol Regulatory Element Binding Transcription Factor 1 (SREBF1) in the regulation of astroglial commitment of hiPSC-NSCs.

Overall, these comprehensive *in vitro* and *in vivo* analyses provide transcriptional and epigenetic reference datasets to define the maturation stage of NSCs derived from different hiPSC sources, and to clarify the safety profile of hiPSC-NSCs, supporting their continuing development as an alternative to somatic hfNSCs in treating neurodegenerative and demyelinating disorders.

## INTRODUCTION

Neural stem cells (NSCs) are promising advanced therapy medicinal products for cell therapies of neurodegenerative disorders^1–4^. NSCs engraft and persist in the brain perivascular stem cell niches, migrate to lesioned areas, and exert neuroprotective and immunomodulatory functions by reducing inflammation and enhancing endogenous repair mechanisms^5–9^. Recent evidence suggests that neurodegenerative disorders are associated with neuroinflammation and multicellular dysfunctions^10–13^, and the peculiar bystander effects and multipotency that distinguish NSCs from committed progenitors and terminally differentiated cells might translate into improved clinical outcomes in this context. Indeed, NSCs have been exploited in pre-clinical studies to treat both acute and chronic neurodegenerative disorders^14–23^. The implementation of Good Manufacturing Practice (GMP)-grade processes for the isolation/enrichment, expansion, and cryopreservation of somatic NSCs from fetal tissues^24^ favored the clinical translation of NSC-based therapies for the treatment of Parkinson’s disease (NCT03128450), ischemic stroke (NCT03296618), spinal cord injury (NCT02163876, NCT01725880, NCT01321333^25^), multiple sclerosis (NCT03282760, NCT03269071^26^), amyotrophic lateral sclerosis (NTC01640067)^27^, neuronal ceroid lipofuscinosis (NCT00337636, NCT01238315^28^), Pelizaeus-Merzbacher disease (NCT01005004, NCT01391637), and cerebral palsy (NCT03005249). Several of these clinical trials have documented safety and some evidence of stabilization of the clinical phenotypes^25–28^. Still, cell therapy based on somatic NSCs presents some challenges, including the production of large amounts of donor cells (in the order of hundreds of millions) under GMP conditions and the immunosuppressive regimens required due to the allogeneic transplant setting. The generation of NSCs from human induced pluripotent stem cells (hiPSCs) might overcome these difficulties, as hiPSCs could be (i) generated from easily accessible cell sources, expanded and differentiated for large-scale GMP production of transplantable cells^29^; (ii) manipulated to create ‘off-the-shelf’ donor cells^30,31^ or collected in HLA-typed hiPSC banks^32^. Additionally, hiPSCs can be genetically modified through lentiviral vector (LV)-mediated gene addition strategies or site-specific gene editing^33–35^, clonally selected based on LV integration profile and genome integrity, and differentiated into transplantable hiPSC-derived NSCs (hiPSC-NSCs), thus potentially limiting genotoxicity in *ex vivo* gene therapy approaches.

While hiPSC-derived neural stem/progenitor cell populations at different stages of commitment – neuroepithelial (NE) cells, radial glia (RG), and NSCs – may display self-renewal and multipotency *in vitro*, their maturation stage might influence their behavior upon transplantation and impact the effectiveness of cell therapies. In the context of demyelinating disorders, intracerebral transplantation of hiPSC-derived NE cells resulted in limited distribution and myelinogenic potential, whereas more committed hiPSC-NSCs sharing phenotypic and functional identity with clinically relevant somatic human fetal NSCs (hfNSCs) displayed an enhanced rostro-caudal migration in white matter regions and gave rise to myelinating oligodendrocytes^33,34,36^. Single-cell RNA-seq (scRNA-seq) analyses revealed intrinsic heterogeneity in NE cell populations, which are enriched in neuronal and glial progenitors with few cell clusters sharing transcriptional identity with somatic hfNSCs^37^. A more in-depth epigenetic, transcriptional, and functional characterization of hiPSC-derived NSCs could define the relevance of generating hiPSC-NSCs with a “bona fide” hfNSC transcriptional signature, unravel the impact of cell heterogeneity, and predict the tumorigenic risk associated with the reactivation of pluripotency programs^38,39^ in view of the clinical exploitation of hiPSC-NSCs in cell-based therapeutic approaches.

Here, we combined multi-omics technologies (bulk RNA-seq, ChIP-seq, and scRNA-seq) and long-term transplantation studies to define the cell identity/heterogeneity and the safety of hiPSC-NSC populations that we previously characterized as phenotypically and functionally similar to hfNSCs^34^.

## RESULTS

### hiPSC-NSCs display a radial glia-like transcriptional profile

We previously validated the genomic stability and pluripotency of hiPSC clones and their ability to differentiate into hiPSC-NSCs that share functional and phenotypical similarity with clinically relevant hfNSCs. Here, by applying the same differentiation protocol^34^ (Suppl. Fig. 1A), we confirmed that hiPSC-NSCs acquire the expression of NSC markers (PAX6, ROBO2) and downregulate pluripotency genes (OCT4, NANOG, LIN28, SSEA4) at levels similar to an hfNSC line that we have previously functionally characterized and used in cell therapy approaches^40^ (Suppl. Fig. 1B-C).

To unravel the transcriptional dynamics underlying the hiPSC-to-NSC differentiation, we performed RNA-seq analysis on four hiPSC-NSC lines and their parental hiPSC clones (Suppl. Table 1). Supervised analysis of RNA-seq datasets identified 6,030 differentially expressed genes (DEGs) between hiPSC-NSCs and hiPSCs samples (Fig. 1A; Suppl. File 1). Euclidean distance plotting highlighted that samples grouped into three separate clusters according to their different origins, with minimal donor-related differences. hiPSC-NSCs grouped more closely with hfNSCs than with parental hiPSCs, demonstrating the consistency and robustness of the reprogramming and neural differentiation protocols and suggesting that hiPSCs globally acquired an NSC-like transcriptional landscape upon neural commitment (Fig. 1B). Gene ontology (GO) analyses of upregulated genes in hiPSC-NSCs vs. hiPSCs identified biological processes and pathways involved in central nervous system (CNS) development, neurogenesis, and neuronal functions, including master regulators of neural commitment/neurogenesis (PAX6, NEUROD1, POU3F2, FOXN4, MEIS1, FOXA1) and genes encoding voltage-gated ion channel subunits (Fig. 1C-D; Suppl. File 1). Meanwhile, processes and pathways regulating the cell cycle, embryonic development, and tissue morphogenesis were downregulated in hiPSC-NSCs (Fig. 1C), which also showed a robust downregulation of the pluripotency network (e.g., POU5F1, NANOG, MYC), mesodermal (e.g. Brachyury (T), GSC) and endodermal markers (e.g. EOMES, SOX17, AFP), and genes involved in embryonic development (including TBX and HOX genes) (Fig. 1D). Notably, genes used as pluripotent-specific markers to monitor residual iPSC contaminants (VRTN, ZSCAN10, LINC00678, L1TD1, and ESRG)^41^ were also strongly downregulated upon neural commitment (Fig. 1A).

**Figure 1.**
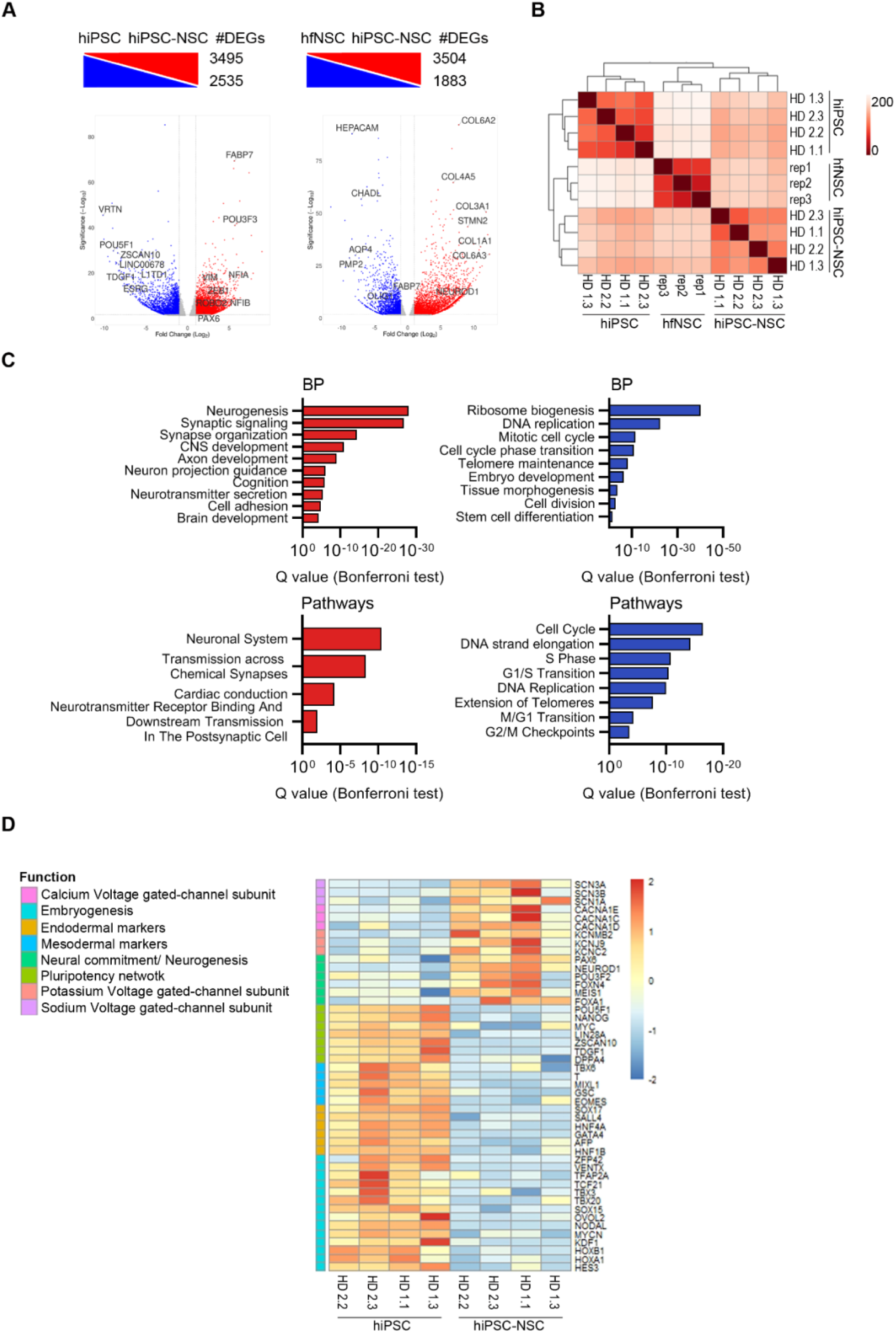
hfNSC-like transcriptional changes during neural commitment distinguish hiPSC-NSCs from parental hiPSCs. **A)** Differentially expressed genes (DEGs) upregulated (red) and downregulated (blue) in hiPSC-NSCs vs. hiPSCs (left) and hiPSC-NSCs vs. hfNSCs (right) (log_2_ fold change ± 1, adjusted *p*-value < 0.05). The total number of DEGs in each comparative analysis is shown. Specific genes with relevant functions in the different cell populations are labeled. **B)** Heatmap of sample-to-sample distance among RNA-seq samples (hiPSCs: clones HD 1.1, HD 1.3, HD 2.2, and HD 2.3; hiPSC-NSC: clones HD 1.1, HD 1.3, HD 2.2, and HD 2.3; hfNSCs, three biological replicates harvested at different passages (p19, p23, p25). **C)** Bar plots of gene ontology enrichment analysis in the comparison hiPSC-NSCs vs. hiPSCs. Enriched biological processes (BP, top plots) and pathways (bottom plots) upregulated (red bars) or downregulated (blue bars) in hiPSC-NSCs are shown. **D)** Heatmap showing the expression levels in hiPSCs and hiPSC-NSCs of genes involved in pluripotency, embryogenesis, mesodermal and endodermal differentiation, neural commitment, and synaptic signaling. Color scale indicates the normalized expression levels of these genes in each sample (blue, low; red, high).

Since transcription factors (TFs) play pivotal roles in cell identity specification and differentiation/maturation processes, we identified the TFs that were up- or downregulated during hiPSC differentiation by integrating RNA-seq datasets with published lists of human TFs^42,43^ (Suppl. File 1). GO analyses of TFs upregulated in hiPSC-NSCs revealed their involvement in cell fate commitment and differentiation processes, with an enrichment of GO terms related to forebrain development, neurogenesis, and gliogenesis (Fig. 2A). Ingenuity Pathway Analysis (IPA) identified upstream regulators driving neural commitment/specification (e.g. PAX6, NEUROD1, OTX2)^44–46^, NSC self-renewal/maintenance (e.g. NUPR1, ZEB1)^47,48^, FOXO3-regulated glucose and glutamine metabolism and autophagy^49,50^, neuronal/glia differentiation (e.g. CREBBP, ZEB2, KDM5A)^51–53^ and repression of the RB-dependent cell cycle (RBL1, ZBTB17) (Suppl. Table 2). Protein-protein interaction enrichment analyses of TFs upregulated in hiPSC-NSCs revealed that genes at hubs and nodes of the regulatory network regulate NSC proliferation/survival (e.g. JUN, CREB5)^54,55^, neuronal/glia commitment/survival (PPARA, RXRB, RUNX2, ATF2)^56–59^, hormone signals (NR3C1, N3RC2, NPAS2, NCOA1)^60,61^ and interferon-JAK-STAT pathways (IRF2, STAT2, STAT3, STAT4)^62–64^ (Suppl. Fig. 1D). Of note, most of these hub TFs were upregulated in the later stages of hiPSC-to-NSC differentiation and NSC expansion, suggesting their role in NSC maintenance/proliferation (Suppl. Fig. 2A).

**Figure 2.**
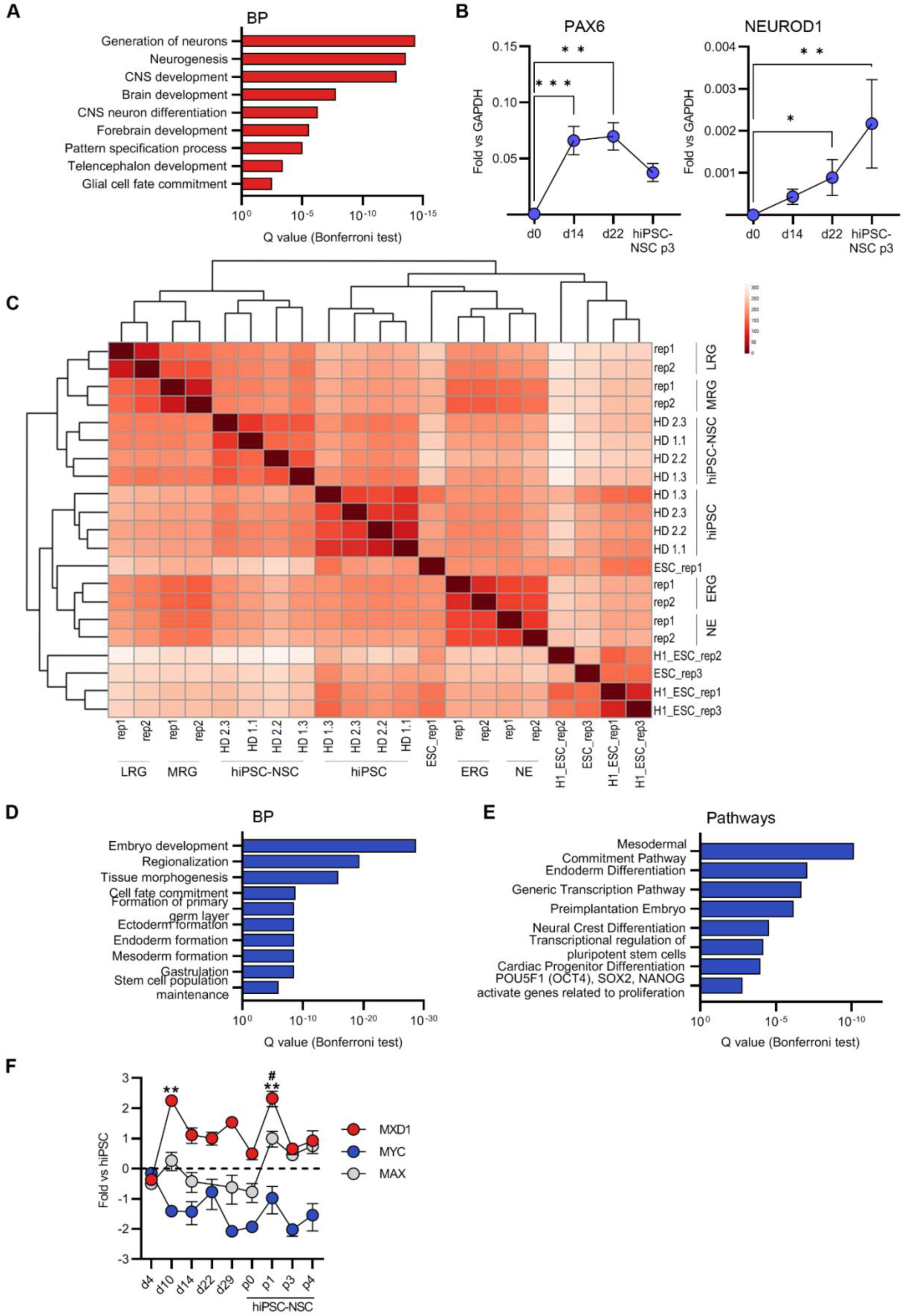
Comparative RNA-seq analyses reveal RG-like transcriptional patterns during hiPSC-to-NSC differentiation. **A)** Gene ontology enrichment analysis of transcription factors upregulated in hiPSC-NSCs relative to parental hiPSCs (log_2_ fold change > 1, adjusted *p*-value < 0.05). Bar plot shows selected biological processes. **B)** qRT-PCR analysis of PAX6 and NEUROD1 expression during hiPSC neural differentiation [timepoints analyzed: hiPSCs (day 0); rosette-like formations (day 14); NE maturation (day 22); and hiPSC-NSCs at passage 3]. Expression levels are normalized on the housekeeping gene GAPDH. Each dot represents the mean ± SEM of 3-4 biological replicates. One-way ANOVA followed by Dunn’s multiple comparison test: *, *p* < 0.05; **, *p* < 0.01; ***, *p* < 0.001. **C)** Heatmap of sample-to-sample distance among RNA-seq samples [hiPSCs: clones HD 1.1, HD 1.3, HD 2.2, and HD 2.3; hiPSC-NSCs: clones HD 1.1, HD 1.3, HD 2.2, and HD 2.3; Embryonic Stem Cells (ESCs): 3 biological replicates (line H1) from^179,180^ and 2 biological replicates (ESC) from^67^; ESC-derived neuroepithelial cells (NE) and early (ERG), middle (MRG), and late (LRG) radial glia cells^67^: 2 replicates]. **D-E)** Gene ontology enrichment analysis of transcription factors downregulated in hiPSC-NSCs relative to parental hiPSCs (log_2_ fold change > 1, adjusted *p*-value < 0.05). Bar plot shows selected biological processes (C) and pathways (D). **F)** qRT-PCR analysis of Myc, Max, and Mxd1 expression during hiPSC-to-hiPSC-NSC transition (timepoints analyzed: Embryoid bodies (day 4), early and late rosette-like formations (days 10 and 14), hiPSC-NSC maturation (days 22 and 29), and hiPSC-NSC at passages 0,1,3 and 4). Expression levels are normalized on the housekeeping gene GAPDH. Each dot represents the mean ± SEM of 3 biological replicates. One-way ANOVA followed by Dunn’s multiple comparison test: **, *p* < 0.01 (vs. d4); #, *p* < 0.05 (vs. hiPSC-NSC p0).

A time-course analysis along hiPSC-to-NSC differentiation revealed that PAX6 and NEUROD1 were upregulated in the early phases of the process, with NEUROD1 being activated at later timepoints likely because of PAX6-mediated transcriptional control of its expression^65,66^ (Fig. 2B). While NEUROD1 expression remained stable during NSC expansion (up to three passages – see Suppl. Fig. 1A), PAX6 was downregulated, suggesting the transition of hiPSC-NSCs from PAX6^high^ NE-like cells to PAX6^low^ RG^67^ (Fig. 2B). Similarly, we observed an increased expression of TFs regulating neural specification/commitment of pluripotent cells (FOXA1, MEIS1)^68–70^ in the early stages of differentiation, whereas TFs driving RG self-renewal/survival (ZNF711, ZEB1, HES1, NUPR1)^47,48,71,72^ were upregulated in hiPSC-NSCs (Suppl. Fig. 2A). Interestingly, TFs driving NSC commitment toward the neuronal lineage were upregulated at both early (POU3F2, RXRB, GRHL1)^56,73,74^ and late (e.g. FOXG1)^75^ timepoints in hiPSC differentiation (Suppl. Fig. 2A). To better characterize the maturation stage of hiPSC-NSCs, we compared our RNA-seq datasets with publicly available transcriptional data on NE cells and RG derived from human embryonic stem cells (ESCs)^67^. Unsupervised analyses of the Euclidean distance among samples highlighted the higher degree of similarity between the transcriptional profiles of hiPSC-NSCs and middle and late RG (MRG, LRG), whereas cells in the earliest steps of differentiation, i.e. NE cells and early RG (ERG), clustered separately (Fig. 2C). In hiPSC-NSCs, the expression levels of core and co-binding factors that functionally regulate the later stages of ESC neural commitment^67^ are expressed at an intermediate level between MRG and LRG with high inter-sample variability, suggesting that hiPSC-NSCs include heterogeneous RG populations at different maturation stages (Suppl. Fig. 2B; Suppl. File 1). Indeed, hiPSC-NSCs showed downregulation of gliogenic genes (OLIG2, NFIA, and NFIB)^76^ and biological processes regulating gliogenesis, and upregulation of neurogenic factors (NEUROD4, NEUROG2) in comparison to fully mature ESC-derived LRG that displays an higher gliogenic potential^67^ (Suppl. Fig. 2B-C; Suppl. File 1).

On the other hand, GO analyses of TFs significantly downregulated during hiPSC differentiation revealed roles in the control of the pluripotency network, embryonic development, and cardiac and neural crest differentiation (Fig. 2D-E; Suppl. File 1). We confirmed these findings by IPA on upstream regulators (Suppl. Table 2), identifying downregulation of pathways downstream to master pluripotency regulators (NANOG, POU5F1) and TFs involved in mesenchymal-to-epithelial transition (TFAP2C)^77^, generation of non-CNS cells (HNF1B)^78^ and cell proliferation (CCN1D, E2F proteins), including the Myc pathways. The activation in hiPSC-NSCs of pathways downstream to MXD1 (Suppl. Table 2; Fig. 2F), which competes with MYC for binding to the MAX co-factor, may repress Myc pathways and favor cell growth arrest and differentiation^79^. Of note, the inhibitory effect of MXD1 on MYC might occur during the initial stages of hiPSC neuroectodermal commitment, since MYC downregulation was concomitant with the upregulation of MXD1 at the earliest timepoints of hiPSC differentiation (Fig. 2F).

Overall, these comparative analyses of RNA-seq data showed that the hiPSC-to-NSC transition involved an initial upregulation of TFs driving neural commitment, followed by the activation of pathways regulating NSC maintenance and multipotency in newly generated hiPSC-NSCs. Human iPSC-NSCs downregulated the transcriptional networks associated with hiPSC pluripotency and commitment toward non-neural cell lineages, acquiring an RG-like transcriptional landscape.

### Enhancers and super-enhancers are the main drivers of the hiPSC-to-NSC transition

Next, we used chromatin immunoprecipitation and sequencing (ChIP-seq) to further characterize hiPSC-to-NSC transition and NSC identity by detecting genomic regions enriched for the histone mark H3K27ac, an epigenetic mark associated with transcription activation. The immunoprecipitated (IP) output of each sample was validated by qPCR against cell-specific regulatory regions of hiPSC and NSC markers (Suppl. Fig. 3A). We identified similar numbers of H3K27ac^+^ promoters in hiPSCs, hiPSC-NSCs and hfNSCs, while the numbers of enhancers and SEs were more variable (Suppl. Fig. 3B). We confirmed that the genes close to and potentially contacted by enhancers and SEs had overall higher expression levels compared to the global expression levels (Suppl. Fig. 3C). Of note, we observed cell-specific differential H3K27ac enrichment in regulatory regions of hiPSC markers (POU5F1, LIN28A) and NSC genes (PAX6, POU3F2) (Suppl. Fig. 3D). The cell-specificity of selected regions was validated by demonstrating that the global expression levels of genes close to cell-specific enhancers and SEs in the cell population of interest was higher compared to their expression levels in other cell populations^80^ (Suppl. Fig. 3E).

When comparing ChIP datasets, we observed that enhancer and SE usage dramatically changed during hiPSC-to-NSC transition, whereas most of the identified H3K27ac^+^ promoters were active in both cell populations (Fig. 3A; Suppl. File 2). These findings suggest the major role of these regulatory regions in driving neural commitment of pluripotent cells. Indeed, by integrating ChIP-seq and RNA-seq datasets and performing GO and IPA analyses, we noted that hiPSC-NSC-specific enhancers and SEs are close to and potentially contacted genes involved in neurogenesis (including EHMT1, CTNNB1, NF-κB1, and TEAD2)^81–84^, synapse organization, neurotransmission, and gliogenesis (Fig. 3B; Suppl. Fig. 3F; Suppl. Table 3; Suppl. File 2). Indeed, these enhancers were enriched in consensus binding sites of TFs involved in neural specification (MEIS1, NEUROD1)^85,86^, RG maintenance, and multipotency (SOX6, RFX2, OLIG2)^87–89^(Fig. 3C; Suppl. File 2). Cell-specific SEs are close to and potentially contacted genes involved in the specification of cholinergic and serotoninergic neurons (Fig. 3B), which could be generated in cultures upon spontaneous differentiation of hiPSC-NSC^34^.

**Figure 3.**
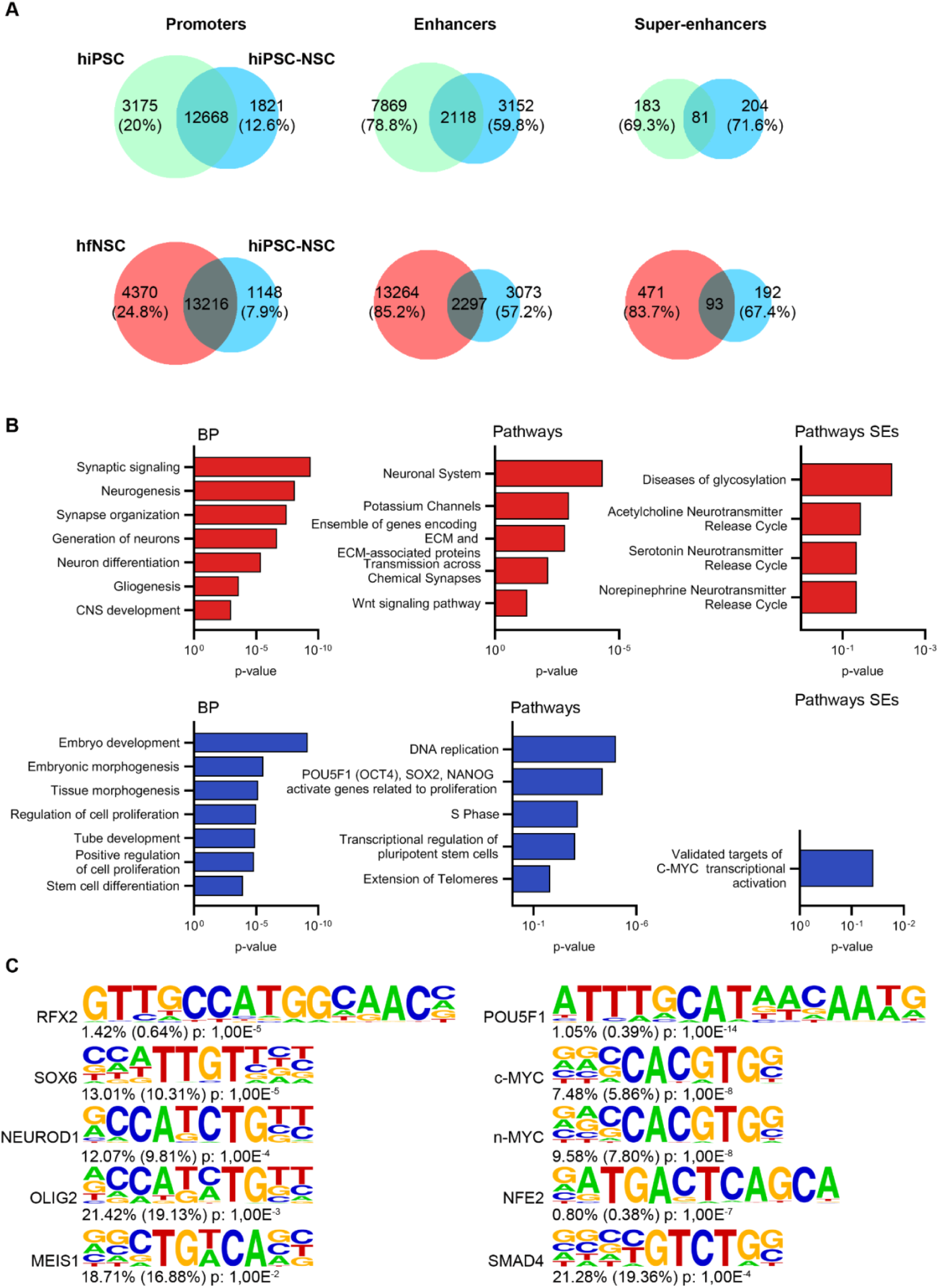
H3K27Ac+ regulatory regions drive the hiPSC-to-hiPSC-NSC transition. **A)** Venn diagrams show the number of cell-specific and shared H3K27ac^+^ promoters, enhancers, and super-enhancers (SEs) between hiPSCs vs. hiPSC-NSCs and hfNSCs vs. hiPSC-NSCs. Percentage indicates the cell-specific regulatory regions in each cell population. **B)** Gene ontology enrichment analysis of up- and downregulated genes (log_2_ fold change ± 1.5 for enhancers and ± 1 for SEs, adjusted *p*-value < 0.05) close to and potentially contacted by cell-specific enhancers and SEs (400 kb window) in hiPSC-NSCs (red bars) vs. hiPSCs (blue bars). Bar plots show the biological processes (BP, left plots) and pathways (middle plots) enriched in enhancers and the pathways (right plots) enriched in SEs for both cell populations. **C)** HOMER analysis of putative transcription factor binding sites (TFBS) detected in hiPSC-NSC-specific enhancers (left) and hiPSC-specific enhancers (right). For each TFBS motif, the percentage of enhancers containing the putative TFBS and the corresponding *p*-value are shown. The background sequences in the whole genome are indicated in parentheses.

Meanwhile, hiPSC-specific enhancers and SEs are close to and potentially contacted genes involved in embryogenesis, the pluripotency network, and MYC pathways (Fig. 3B; Suppl. File 2). TF binding sites (Fig. 3C; Suppl. File 2) and IPA analyses (Suppl. Table 3; Suppl. Fig. 3F) showed that hiPSC-specific enhancers and SEs regulate pluripotency genes (NANOG, POU5F1, MYC^90^) and TFs involved in the cell cycle (E2F1, MYBL2)^91,92^, non-neural tissue specification (MYOCD; GATA4)^93,94^, DNA damage response (NER, BER, ATM signaling), and tissue-specific cancer.

Overall, these data indicate that enhancers and SEs are the main drivers of the transcriptional changes occurring during the hiPSC-to-NSC transition, with hiPSC-NSC-specific regulatory regions positively regulating the main traits of RG identity, including cell specification and multipotency. Of note, hiPSC-to-NSC commitment results in the inactivation of enhancers regulated by the pluripotency core of transcription factors, reinforcing the switch-off of the hiPSC pluripotency network at both regulatory and transcriptional levels.

### hiPSC-NSCs and hfNSCs include RG at different stages of maturation

To better highlight similarities and differences among hiPSC-NSCs and hfNSCs we compared their transcriptional and epigenetic landscapes (RNA-seq and ChIP-seq datasets). Both NSC populations shared common H3K27ac^+^ promoters (Fig. 3A; Suppl. File 2) and pathways related to the cell cycle and metabolism as well as NGF and CXCR4 signaling, involved in NSC survival and migration^95,96^ (Suppl. Fig. 4A-B; Suppl. File 1-2). Enhancer and SE were differentially activated in hiPSC-NSCs as compared to hfNSCs (Fig. 3A; Suppl. File 2) and we detected upregulation of transcripts and genes involved in extracellular matrix organization, likely explained by the adherent culture conditions (Fig. 4A-B; Suppl. File 1-2). In contrast, hfNSCs growing as floating neurospheres upregulated cell-cell adhesion processes and pathways involved in glycosaminoglycans and chondroitin sulfate, signaling molecules of the neurogenic niche^97^ (Fig. 4A-B). These results highlight the impact of culture conditions on the overall transcriptional and epigenetic profiles of these cells.

**Figure 4.**
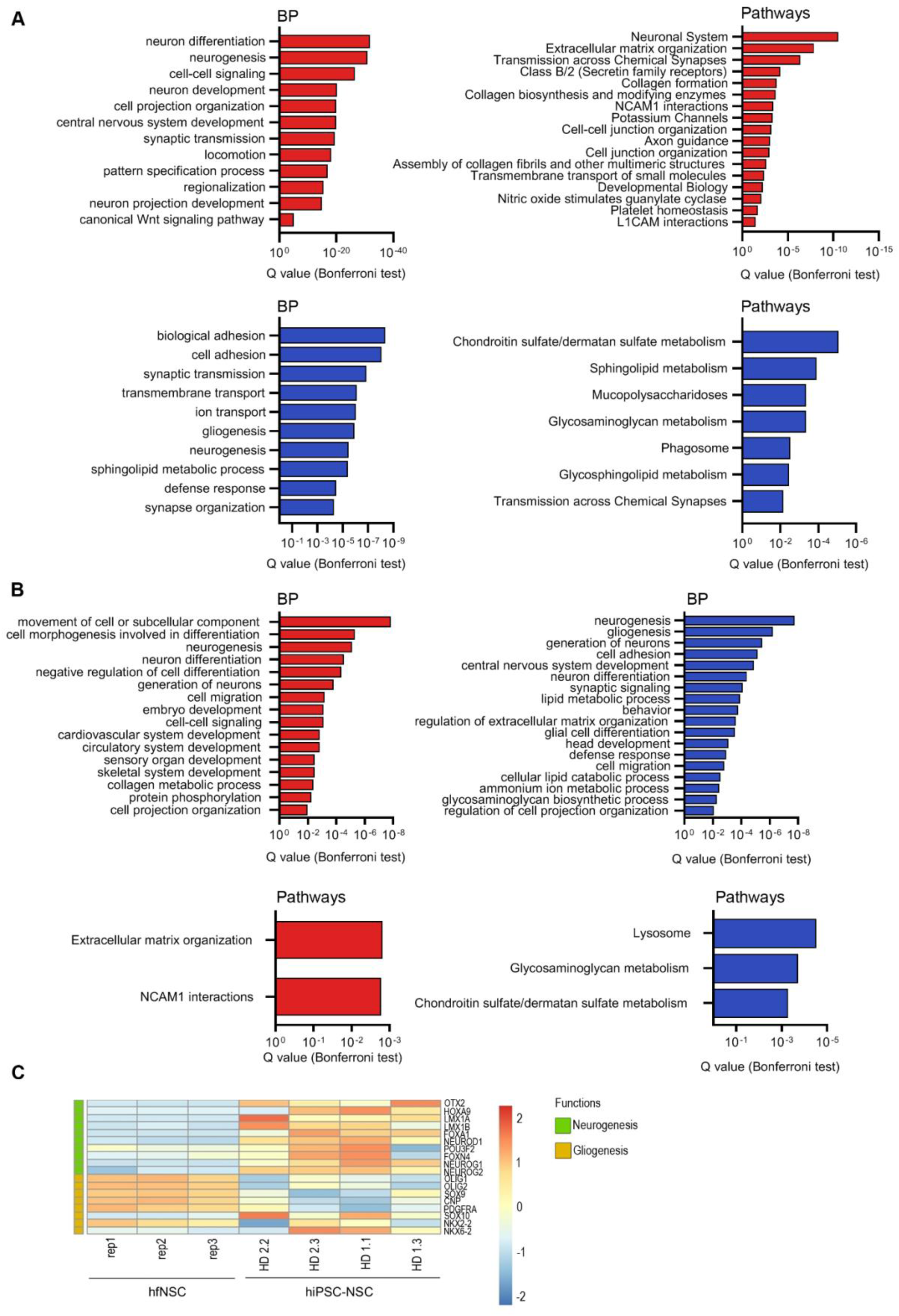
Transcriptional and epigenetic profiles reveal differences in differentiation potential in hiPSC-NSC and hfNSC. **A)** Gene ontology enrichment analysis of RNA-seq data in the comparison of hiPSC-NSCs vs. hfNSCs (log_2_ fold change ± 1, adjusted *p*-value < 0.05). Bar plots represent the biological processes (BP, left plots) and pathways (right plots) upregulated (red bars) or downregulated (blue bars) in hiPSC-NSCs as compared to hfNSC. **B)** Gene ontology enrichment analysis resulting from the integration of ChIP-seq and RNA-seq datasets in the comparison of hiPSC-NSCs vs. hfNSCs. Bar plots show BP (upper plots) and pathways (bottom plots) of upregulated genes in hiPSC-NSCs (red bars) or upregulated genes in hfNSCs (blue bars) (log_2_ fold change ± 1,5, adjusted *p*-value < 0.05) close to cell-specific enhancers (100 kb window). **C)** Heatmap showing the expression levels of neurogenesis-associated genes and oligodendrocyte markers in hiPSC-NSCs vs. hfNSCs. Color scale indicates the average expression levels of these genes in each cluster (blue, low; red, high).

Additionally, hiPSC-NSCs are characterized by higher activation of pathways downstream TFs regulating brain development (ARNT2, OTX2, POU3F2)^98,99^, NE-to-RG transition (NOTCH3, LMX1a)^100,101^, pattern specification of ventral neural progenitors (HOXA9, LMX1a, LMX1b, OTX2)^99,102^, and quiescence/maintenance of the NSC pool (FOXO1, KLF4)^103^(Fig. 4B; Suppl. Table 4; Suppl. Fig. 4C; Suppl. File 1-2). The increased expression of proteins and TFs involved in the WNT/β-catenin (e.g. CTNNB1, LEF1)^98^ and Hedgehog (e.g. GLI1) pathways and in BMP and SMAD signaling (SMAD4, SMAD3, SOX11)^98,104^ could be associated with the propensity of hiPSC-NSCs to generate neurons (Fig. 4A-C; Suppl. Table 4). On the other hand, cell-specific enhancers/SEs and genes regulating gliogenesis (including the pathways downstream from SOX3 and PRMD8)^60,105^ were more highly expressed in hfNSCs (Fig. 4A-C; Suppl. Table 4; Suppl. Fig. 4C). Notably, the residual activation of enhancers regulating genes driving embryonic and non-CNS development (Fig. 4B) is associated with pathways commonly activated during the commitment of pluripotent stem cells toward the neural lineage. Indeed, downstream analysis revealed that 58.74% of these genes were shared among GO terms describing the development of CNS and non-neural tissues (Suppl. Fig. 4D).

To dissect the intra- and inter-sample heterogeneity and define the cell identity and safety of the different NSC populations we performed single-cell RNA-seq (scRNA-seq) on hiPSC-NSCs and hfNSCs. After filtering and normalization, our dataset included the transcriptomes of 1,000-3,000 cells/sample. By using gene RNA-seq and scRNA-seq signatures from hiPSC- or ESC-derived neural cells at different stages of maturation and from neuronal/glia progenitors^37,61,67,106,107^, we confirmed a higher expression of ERG and MRG markers in hiPSC-NSCs as compared to hfNSCs, which are mainly composed of cells with transcriptional signatures resembling LRG and oligodendroglial precursors (Suppl. Fig. 5A). By integrating our datasets with scRNA-seq gene signatures of RG from the human fetal brain^108^, we observed a higher transcriptional identity of hiPSC-NSC with immature ventricular RG (vRG), while hfNSC showed similarity with mature outer RG (oRG) and pre-oligodendrocyte progenitors. (Suppl. Fig. 5B).

Uniform Manifold Approximation and Projection (UMAP) representation revealed intra-sample heterogeneity in both hiPSC-NSC and hfNSC samples (Fig. 5A), leading to the annotation of 12 clusters characterized by different transcriptomic profiles (Fig. 5B; Suppl. File 3). Pseudotime analyses of UMAP clusters identified a trajectory describing the progressive transition from immature to mature RG and the acquisition of a neuronal progenitor-like signature (Fig. 5C, Suppl. Fig. 5C). Indeed, the expression of PAX6 and ERG markers was higher in Cluster 1 (PAX6^high^ / OTX2^low^ RG) than in Cluster 2 (PAX6^low^ / OTX2^high^ RG), which still maintained the transcriptional signature of MRG (Fig. 5D-E). Additionally, Cluster 2 showed a higher activation of N-glycan biosynthesis, associated with the neurogenesis/astrogenesis switch in NSCs^109^, and glycolysis, which is typically activated in neural stem cells to provide energy for maintenance of stemness^110^ (Fig. 5F; Suppl. File 3). Along the pseudotime trajectory, Cluster 3 (SLC1A3^high^ RG), composed of a mix of hiPSC-NSCs and hfNSCs, was characterized by high expression of LRG markers, activation of cAMP and calcium signaling, and positive regulation of the PPAR pathway and cholesterol metabolism linked to oligodendrogenesis^111–115^ (Fig. 5D-F). Cluster 5 (GBX2^high^ RG) might represent RG that progressively acquired a neuronal progenitor phenotype, with higher expression of known neuronal precursor markers in Cluster 7 (neuronal precursors) (Fig. 5D-E). Cluster 7 was characterized by activation of pathways regulating RAS and ERBB signaling involved in the proliferation, survival, differentiation, and migration of neuronal progenitors^116–120^, axon guidance, synaptic vesicles, and glutamatergic and GABAergic synapses (Fig. 5F). Moreover, cells within cluster 7 have a transcriptional profile similar to neuronal cells of the human fetal brain ^108^ since we identified transcripts expressed in intermediate progenitors (IP) and mature excitatory/inhibitory neurons along the pseudotime trajectory (Suppl. Fig. 5B). Of note, Clusters 4 (cycling SLC1A3^+^ RG) and 6 (cycling GBX2^high^ RG) displayed transcriptional profiles similar to Clusters 3 and 5, respectively, with higher activation of cell cycle genes (Fig. 5D-F). Additionally, we have identified a cluster mainly composed of hfNSCs expressing typical LRG genes (Cluster 11).

**Figure 5.**
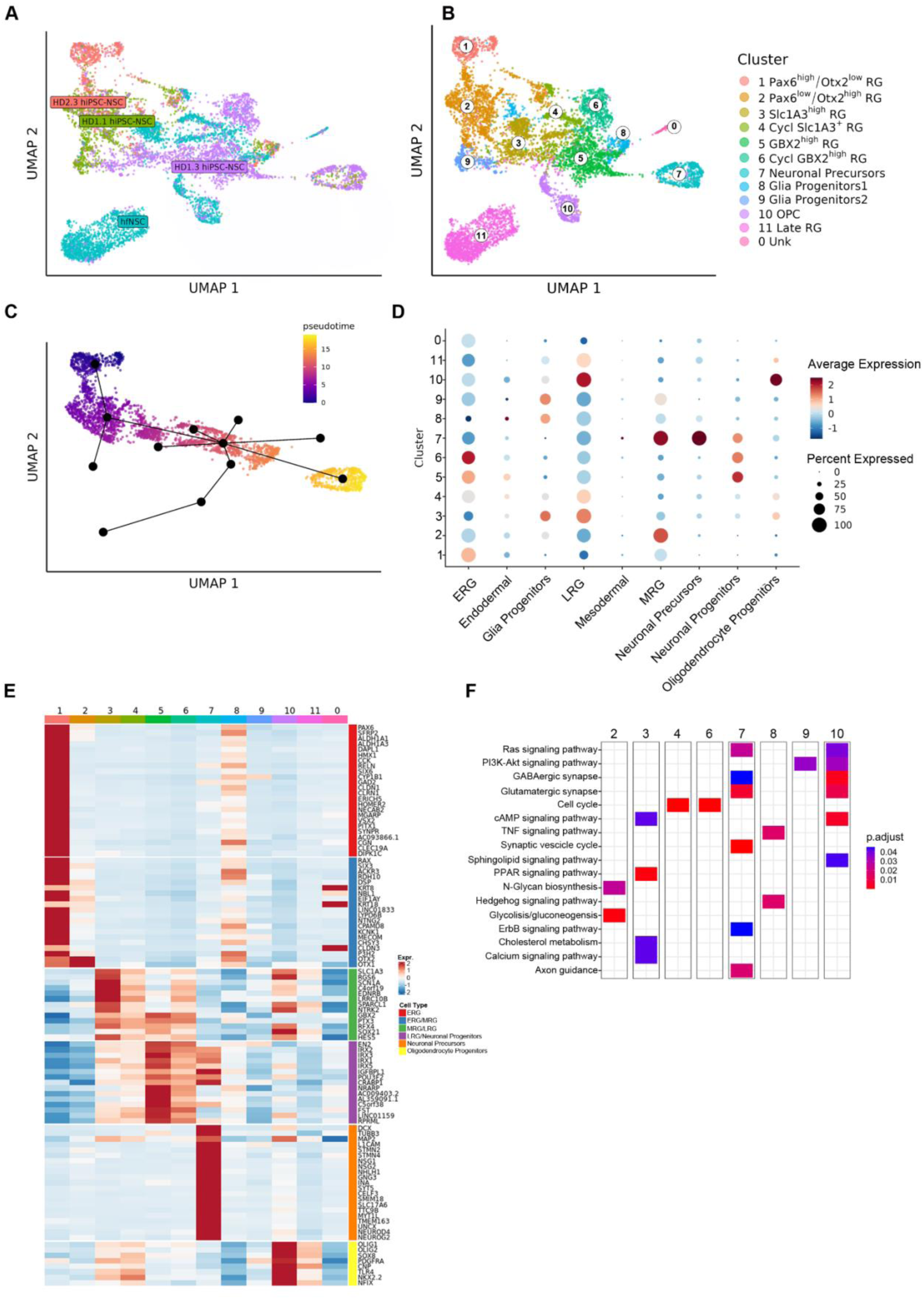
Heterogeneity in hiPSC-NSC cell composition reflects a different degree of RG maturation. **A)** UMAP plot showing the distribution of scRNA-seq transcriptomes (∼1,000-3,000 cells/sample) of three hiPSC-NSC clones [HD 1.1 (green dots), HD 1.3 (purple dots), and HD 2.3 (red dots)] and hfNSC (blue dots); resolution 0.6. **B)** UMAP plot showing numbers and labels of the different clusters identified in scRNA-seq analyses (resolution 0.6). Each cluster was annotated based on published NSC transcriptome datasets and the expression of cell-specific markers. **C)** Pseudotime analysis showing the transcriptional trajectory that describes the progressive transition from Cluster 1 to Cluster 7. **D)** Dot plot showing the signature annotation of each cluster based on published datasets as described in (B). Dot size indicates the percentage of signature-specific genes expressed in each cluster. Average expression levels of these genes in each cluster are depicted according to the color scale shown (blue, low; red, high). **E)** Heatmap showing the expression levels of signature-specific markers in each cluster. Color scale indicates the average expression levels of these genes in each cluster (blue, low; red, high). **F)** Heatmap showing selected pathways (KEGG database) enriched in the different scRNA-seq clusters. The color scale reports the statistical significance (adjusted *p*-value) of each pathway in the clusters.

We detected higher expression of markers of ESC-derived glia progenitors in the hiPSC-NSC-enriched cluster 8, a subpopulation of TNF- and Hedgehog-responsive glia progenitors (Glia Progenitors 1), and in Cluster 9, characterized by the activation of PI3K-AKT and sphingolipid signaling (Glia Progenitors 2) (Fig. 5D, F). Markers of oligodendrocyte progenitors were upregulated in Cluster 10 (OPC), which was mainly composed of hfNSCs (Fig. 5D-E). Of note, more committed neuronal (Clusters 5, 6 and 7) and oligodendroglial (Cluster 10) subpopulations were characterized by activation of the oxidative phosphorylation pathway, which favors a more efficient energy production to match the needs of the differentiating progeny^110^ (Fig. 5F). Importantly, we defined transcriptional signatures (Fig. 5D) and membrane-bound markers (Suppl. Fig. 5D) to identify hiPSC-derived RG at different stages of maturation and committed neuronal and glia progenitors. Interestingly, the top 50 genes that identify hiPSC-NSC/hfNSC-derived mature RG (Clusters 3-6 and 11) and committed progenitors (clusters 7, 10) were detected in RG and glial/neuronal progenitor cell subpopulations of the human fetal brain (Suppl. Fig. 5E)^108^. Importantly, we did not detect Oct4^+^ cells in hiPSC-derived subpopulations and detected only low residual expression of mesodermal/endodermal markers or genes involved in hiPSC/ESC pluripotency (e.g. CNMD, DPPA4 and L1TD1)^121–123^(Fig. 5D, Suppl. Fig. 5A, F).

Overall, these data highlight the heterogeneity of the hiPSC-NSC and hfNSC populations, which are composed of RG at different maturation stages and display distinct regionalization and gliogenic vs. neuronal potential.

### SREBF1 is involved in the regulation of astroglia commitment

The scRNA-seq analyses showed that genes associated to *de novo* lipogenesis were highly expressed in the hiPSC-NSC-derived glia progenitors (Cluster 8) (Suppl. Fig. 6A), suggesting that this pathway may be involved in glial commitment/differentiation alongside its well-known role in NSC function^124,125^. Interestingly, bulk RNA-seq and ChIP-seq data highlighted hiPSC-NSC-specific SEs driving the expression of genes downstream the Sterol Regulatory Element Binding Transcription Factor 1 (SREBF1), a key transcriptional activator of genes involved in cholesterol biosynthesis and lipid homeostasis^126^ (Suppl. Table 3). A time-course gene expression analysis showed that SREBF1 is upregulated in the late stages of hiPSC-to NSC differentiation and in hiPSC-NSCs (Suppl. Fig. 6B), suggesting its potential role in NSC commitment/maintenance.

To test this hypothesis, we generated SREBF1 knockout (KO) hiPSC lines by co-delivering ribonucleoprotein Cas9 with a pool of sgRNA targeting exon 5 of the SREBF1 gene. We achieved 89% KO efficiency in the bulk hiPSCs, as detected by Synthego ICE and qRT-PCR analyses (Suppl. Fig. 6C). By subcloning Cas9/sgRNA-treated hiPSCs, we isolated three clones with 100% KO efficiency (Suppl. Fig. 6D) that were then differentiated into hiPSC-NSCs. The abrogation of SREBF1 expression in hiPSC-NSCs did not impact cell survival (Suppl. Fig. 6E). SREBF1 KO hiPSC-NSC clones showed similar expression levels of NSC (PAX6, FOXG1, NEUROD1) and neuronal (MAP2, DCX) genes and a 4.5-fold decrease in the expression of the GFAP gene in comparison to Cas9-only control cells (Suppl. Fig. 6F). To investigate the impact of SREBF1 KO on hiPSC-NSC multipotency, we differentiated hiPSC-NSCs into a mixed neuronal/glial cell population using an optimized protocol^34^. SREBF1 KO neuronal/glial cultures showed similar expression levels of neuronal genes (NEUROD1, DCX, β-tubulin III, TH), while the expression of astroglial genes (GFAP and FABP7) was significantly reduced in comparison to control cells (Suppl. Fig. 6G). Overall, these data identify SREBF1 as a key TF favoring the astroglial commitment/differentiation of hiPSC-NSCs, with implications for possible drug-mediated modulation of the gliogenic potential of this population.

### hiPSC-NSCs are transcriptionally divergent from glioblastoma stem cells

The major safety concern in cell therapy with hiPSC-derived cells is the potential residual expression or reactivation of cancerogenic pathways in transplanted cells^127,128^. Our comparison of RNA-seq and ChIP-seq datasets from hiPSC-NSCs and hfNSCs suggested the potential activation of pathways downstream pro-oncogenic TFs^129,130^ in hiPSC-NSCs (Suppl. Table 2 and Supp. Table 4). Since cancer stem cells in glioblastomas (GBM) share common features and niches with NSCs^131^ we investigated the transcriptional similarities/differences of hiPSC-NSCs and hfNSCs in comparison with primary glioblastoma stem cells (GSCs)^132^.

Euclidean distance among samples indicated that the three cell populations clustered separately, with low similarity between the transcriptional profiles of hiPSC-NSCs and GSCs (G523NS) (Fig. 6A). Supervised analysis of RNA-seq datasets identified 8,471 DEGs in hiPSC-NSC vs. GSC samples (Fig. 6B; Suppl. File 1), highlighting a higher degree of transcriptional diversity relative to the comparison between the two neural populations (Fig. 1A). Of note, we detected 6,626 DEGs in hfNSCs vs. GSCs (Fig. 6B; Suppl. File 1), further suggesting a low propensity for tumorigenesis in clinically-relevant NSC populations. The majority of potential oncogenic TFs detected in hiPSC-NSCs (Suppl. Table 4) as well as key TFs and signaling molecules driving GBM reprogramming and GSC growth and development were expressed at significantly lower levels in hiPSC-NSCs vs. GSCs ^130,133–136^(Fig. 6C).

**Figure 6.**
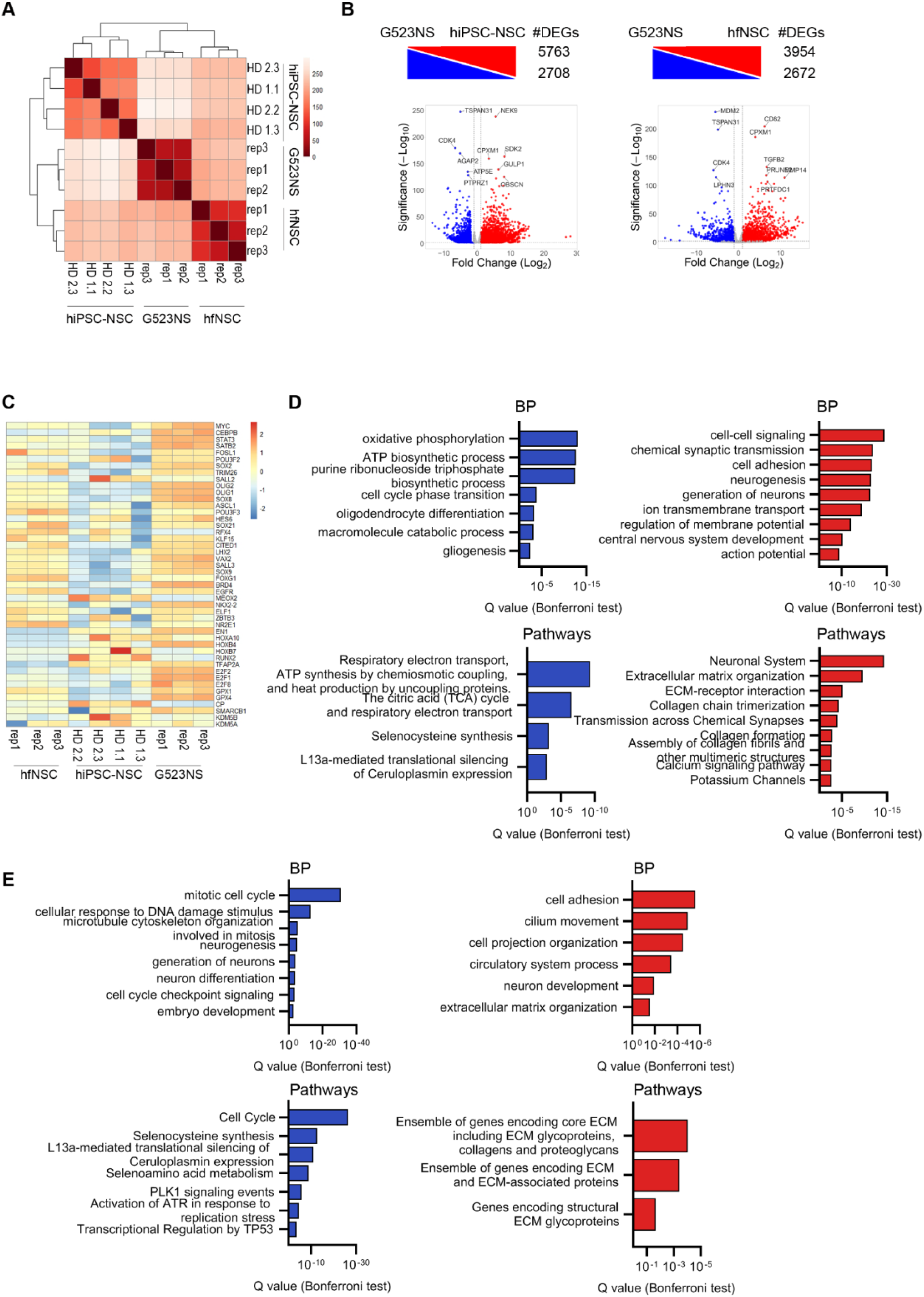
hiPSC-NSCs and hfNSCs are transcriptionally divergent from glioblastoma stem cells. **A)** Heatmap of sample-to-sample distance among RNA-seq samples [hiPSC-NSC: clones HD 1.1, HD 1.3, HD 2.2, and HD 2.3; hfNSCs: three biological replicates harvested at different passages (p19, p23, p25); glioblastoma stem cell (GSCs) line (G523NS): three biological replicates^132^]. **B)** Differentially expressed genes (DEGs) upregulated (red) and downregulated (blue) in hiPSC-NSCs vs. hiPSCs and hiPSC-NSCs vs. hfNSCs (log_2_ fold change ± 1, adjusted *p*-value < 0.05). The total number of DEGs in each comparative analysis is reported. The most up- or downregulated genes with relevant functions in the different cell populations are highlighted. **C)** Heatmap showing the expression of potential GSC markers and pro-oncogenic transcription factors and signaling molecules in hiPSC-NSC, hfNSC, and GSC samples. Color scale indicates the average expression levels of these genes in each cluster (blue, low; red, high). **D-E)** Gene ontology enrichment analysis of upregulated (red bars) and downregulated (blue bars) genes in hiPSC-NSCs vs. GSCs (D) and hfNSCs vs. GSCs (E) (log_2_ fold change ± 1, adjusted *p*-value < 0.05). Bar plots in each panel show selected biological processes (BP, upper plots) and pathways (bottom plots).

GO analyses of biological processes and pathways downregulated in NSC populations vs. GSCs revealed that tumorigenic stem cells showed an increased expression of genes driving the cell cycle (including MYC and E2F TFs); oxidative phosphorylation-dependent ATP biosynthesis, typically detected in cancer vs. somatic stem cells^137^; synthesis of selenoproteins, which play a key role in oxidative stress regulation in glioblastoma (e.g. GPX1 and GPX4^138–143^); and L13a-mediated silencing of ceruloplasmin, an important regulator of iron metabolism which positively correlates with the efficacy of radiotherapy on GBM cells^144^ (Fig. 6C-E; Suppl. File 1). Conversely, both hiPSC-NSCs and hfNSCs upregulated biological processes and pathways involved in collagen organization, which seems to positively correlate with longer median survival in GBM patients^145^, and in neurogenesis or synaptic transmission, which reflects the multipotency of these cell populations relative to GSCs (Fig. 6D-E).

Overall, these data suggest that hiPSC-NSCs are transcriptionally divergent from GSCs, further supporting their safety profile.

### Safe, stable, and long-term engraftment of hiPSC-NSCs upon intracerebral transplantation in mice

Our omics analyses indicated that hiPSC-to-NSC differentiation involves transcriptional and epigenetic silencing of the pluripotency network and cancer-associated pathways and results in a heterogeneous hiPSC-NSC population mainly composed of RG and neuronal progenitors. To finally assess the safety and functionality of these cells *in vivo*, we transplanted hiPSC-NSCs into immunodeficient (Rag2^−/−^/γ-chain^−/−^) neonatal mice (bilateral intracerebroventricular injection; 200,000 cells/brain) and analyzed treated animals at 10 months post-transplant. We employed immunofluorescence analysis with human-specific antibodies (hNuclei and STEM121) and cell lineage-specific markers to evaluate cell engraftment and distribution and the phenotype of transplanted cells engrafted in brain tissues.

We detected variable but robust engraftment of human cells (Fig. 7A-B), which appeared well-integrated into white and gray matter areas, including the olfactory bulbs, striatum, cerebellum, pons, medulla, and cervical spinal cord (Fig. 7A; Suppl. Fig. 7A-B), indicating a widespread rostro-caudal migration from the injection site. We found overall low percentages of human cells expressing the proliferation marker Ki67 (4.4% ± 1.0%). Interestingly, Ki67^+^ human cells were preferentially located within or close to the subventricular zone (SVZ) neurogenic niche (Fig. 7C). The presence of NESTIN^+^ human cells in the SVZ and other brain regions confirmed the long-term persistence of engrafted hiPSC-NSCs retaining an immature phenotype (Fig. 7D-E). The majority of human cells detected in non-SVZ brain regions and the spinal cord expressed markers of astrocytes (S100β) and oligodendrocytes (glutathione S-transferase π, GSTπ) (Fig. 7D-E; Suppl. Fig. 7B).

**Figure 7.**
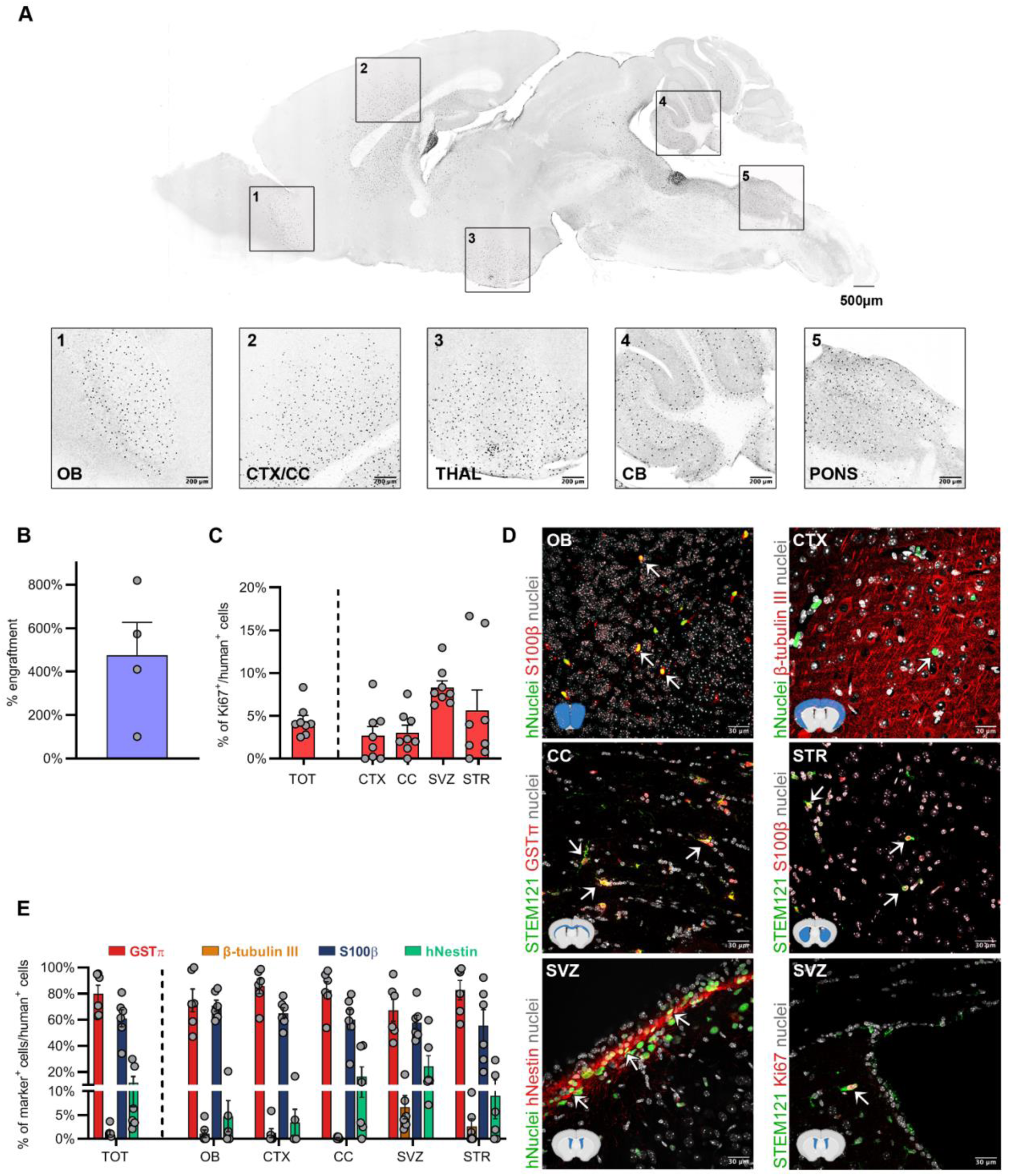
Long-term engraftment of hiPSC-NSCs upon intracerebral transplantation in neonatal mice. **A)** Representative mask of a sagittal brain section generated based on immunofluorescence images, showing the distribution of engrafted HD 2.2 hiPSC-NSCs at 10 months after transplantation. Higher magnifications of specific brain regions are shown: 1. olfactory bulbs (OB); 2. cortex/corpus callosum (CTX/CC); 3. thalamus (THAL); 4. cerebellum (CB); 5. pons/medulla (PONS). Human cells were stained with hNuclei antibody. **B)** Bar plot showing the percentages of engrafted hiPSC-NSCs in the brains of transplanted mice (n = 4). Human cells were stained with hNuclei antibody. The percentage of engraftment was expressed as: [total number of engrafted cells (hNuclei^+^ cells in one series of coronal/sagittal sections × number of series) / total number of transplanted cells] × 100. Each dot represents one mouse. **C)** Bar plot showing the percentage of Ki67^+^ proliferating human cells detected in the entire brain (TOT) and spitted in selected regions (CTX; CC; SVZ: sub-ventricular zone; STR: striatum). Each dot represents one mouse (n=8). **D)** Representative immunofluorescence images of engrafted cells (hNuclei^+^ or STEM121^+^) in different brain regions expressing proliferation (Ki67) and/or cell-specific markers: S100β (astrocytes), GSTπ (oligodendrocytes), β-tubulin III (neurons), hNestin (neural stem cells). Nuclei were counterstained with Hoechst. The imaged brain regions are labeled as in (A,B). Arrows indicate co-localization of immunofluorescence signals. **E)** Bar plot reporting the quantification of the engrafted cells (hNuclei^+^ or STEM121^+^) expressing the cell-specific markers shown in (D). Data reports the quantification performed on the total brain (TOT) and in selected regions (OB, CTX, CC, SVZ, and STR). Percentages are calculated as: (number of cells expressing cell-specific markers) / (number of cells expressing human cell markers) × 100. Each dot represents data collected in one mouse (n=6).

The presence of ≅ 50% and ≅ 80% of S100β^+^ cells and GSTπ^+^ cells, respectively, in the engrafted human cell populations, suggested that these two markers may be co-expressed in subpopulations of glial cells. To verify this hypothesis, we checked for the presence of cells expressing SOX10, a TF expressed by NSCs and a master TF for oligodendrocyte specification whose expression has also been associated with a certain degree of plasticity in terms of astroglial differentiation^146,147^. Double immunofluorescence analyses revealed that the majority of S100β^+^ and GSTπ^+^ cells co-expressed SOX10, suggesting that they may represent a subpopulation of committed glia progenitors (Suppl. Fig. 7C-D). An average of 40% S100β^+^/SOX10^-^ astrocytes and 25% GSTπ^+^/Sox10^-^ oligodendrocytes were detected in grey and white matter regions of transplanted animals, with a higher distribution of hiPSC-NSC-derived oligodendrocytes in the corpus callosum and cortex (Suppl. Fig. 7C-D). We found minor percentages of human-derived β-tubulin III^+^ neurons (2%-20%), which mainly localized in regions close to the injection sites (SVZ and striatum) (Fig. 7D-E; Suppl. Fig. 7B). Importantly, we did not detect any human cells expressing pluripotency markers (NANOG and OCT4), nor did we detect abnormal proliferation or tumor formation in the brains of any hiPSC-NSC-transplanted mice.

## DISCUSSION

The complexity of the transcriptional and epigenetic programs that drive the initial steps of neural commitment in pluripotent stem cells have been thoroughly investigated in ESCs^67,148^ and hiPSCs^149,150^. However, information is lacking on the transcriptional and epigenetic landscape of hiPSC-derived neural cell populations at later stages of maturation. We took advantage of our previously optimized differentiation protocol to generate hiPSC-NSCs with phenotype and function resembling those of somatic hfNSCs isolated from the human fetal brain, which comprise a heterogeneous population of neural stem and progenitor cells with RG-like features displaying long-term self-renewal, proliferation, and multipotency when maintained in appropriate culture conditions^40^.

By investigating the chromatin changes occurring at H3K27ac^+^ active regulatory regions, we observed a dramatic change in the usage of enhancers and SEs during the hiPSC-to-NSC transition, while > 70% of the mapped promoters were shared between hiPSCs and hiPSC-NSCs. This finding confirms that enhancers play a major role in driving the neural commitment of pluripotent cells and in determining the cell identity of ESC- and hiPSC-derived cells^148,151,152^. In hiPSC-NSCs, we observed upregulation of pathways that are usually activated in pluripotent cell-derived neural cells, including CNS development, neurogenesis, and synaptic transmission^67,148–150^. The concomitant upregulation of TF hubs regulating NSC self-renewal/maintenance at later stages of differentiation confirmed the acquisition of the neural fate in hiPSC-NSCs. Epigenetic activation of NSC-specific enhancers and SEs driving neuronal/glia specification and neuronal transmission suggest that active regulatory regions primed hiPSC-NSCs to differentiate into glial cells and neurons, particularly neuronal subpopulations (cholinergic and serotoninergic neurons) that arise *in vitro* upon spontaneous differentiation^34^. Interestingly, transcriptomic data suggested an interdependent regulation of two key players of neural fate specification, PAX6 and NEUROD1. While PAX6 is strongly upregulated until the formation of hiPSC-derived neural rosettes (day 14 of differentiation)^34^, NEUROD1 expression slowly and constantly increases along the differentiation, suggesting that PAX6 triggers NEUROD1 expression in human cells as previously observed in murine somatic NSCs^65^. The later steps of NSC differentiation are characterized by a robust downregulation of PAX6 expression, indicating the acquisition of a PAX6^low^ RG-like phenotype in the majority of hiPSC-NSCs^67^. Indeed, the comparison of transcriptional datasets from hiPSC-NSCs and ESC-derived neural subpopulations^147^ showed the acquisition of a global transcriptomic profile corresponding to an intermediate state between MRG and LRG in hiPSC-NSCs, as confirmed by the expression levels of core and co-binding factors driving the generation of mature ESC-derived RG^67^. Pseudotime analyses on scRNA-seq datasets revealed that hiPSC-NSCs include RG subpopulations in the transition between ERG and LRG. This finding is corroborated by the reduced activation of gliogenic potential, a process typically acquired in the later stage of RG maturation^67^, in hiPSC-NSCs as compared to ESC-derived LRG and somatic hfNSCs. Additionally, hiPSC-NSCs show a transcriptional profile resembling primitive vRG isolated from the human fetal brain^108^. Importantly, we identified novel transcriptional signatures to discriminate immature and mature hiPSC-derived RG, as well as committed progenitors, that can be prospectively applied to characterize hiPSC-derived neural cultures generated with different protocols or to isolate specific subpopulations for targeted cell replacement approaches.

Different pathways regulating metabolic processes play a role in the maturation and differentiation of hiPSC-derived RG. In scRNA-seq and pseudotime analyses, we observed the glycolysis/oxidative phosphorylation metabolic rewiring needed to ensure efficient energy production in committed neuronal and glia cells^110^, activation of the PPAR pathway and cholesterol metabolism linked to oligodendrogenesis^111–115^, and N-glycan biosynthesis/branching promoting astrogenesis switching in NSCs^109^. Interestingly, the transcription factor SREBF1 – a member of the sterol regulatory element-binding proteins (SREBPs) that control lipid metabolic pathways^153^ – is upregulated at the final stages of hiPSC-to-NSC differentiation, and the activation of its downstream pathways in hiPSC-NSCs is associated with cell-specific SEs. We observed that SREBF1 ablation in hiPSCs and NSC progeny does not impact cell survival and NSC-specific gene expression. Since lipid metabolism plays a crucial role in somatic NSC proliferation/quiescence and development through fatty acid synthase (FASN)-dependent *de novo* lipogenesis^125,154–156^, our data suggest the occurrence of compensative pathways to maintain NSC homeostasis (e.g., direct FASN activation by LXRs interacting with its promoter)^157^. While SREBF1 ablation leads to impairment of murine dopaminergic neurons^158^, its role in RG and astrocytes is still unknown. Our findings indicate that the deletion of this transcription factor impacts astroglial commitment and differentiation of hiPSC-NSCs, as illustrated by the decreased expression of typical astrocytic markers in hiPSC-NSC-derived differentiated cultures. Thus, we identified a novel pathway driving gliogenesis in hiPSC-NSCs which can be potentially modulated to increase the generation of hiPSC-derived glial cells, particularly astrocytes.

To be considered as an alternative source for cell therapy approaches, hiPSC-NSCs should share the main clinical safety features of clinically approved hfNSCs, including growth factor-dependent proliferation/survival, comparable engraftment potential in transplanted brains, multipotency, and absence of tumorigenicity. Our epigenetic and transcriptomic analyses show major similarities in the biological processes regulating the cell cycle and metabolic pathways between hiPSC-NSCs and hfNSCs, with a similar activation of enhancers driving NGF-dependent signaling^159,160^ and CXCR4-mediated cell migration^161,162^. In addition to the stage of RG maturation, the transcriptional and epigenetic differences seem to be mainly attributable to the different *in vitro* culture conditions: extracellular matrix interactions with the laminin/poly-L-ornithine coating were upregulated in hiPSC-NSCs (grown in adhesion), whereas chondroitin sulfate metabolism and cell-cell adhesion pathways were activated in floating neurospheres composed of hfNSCs^97^.

Our previous side-by-side functional and phenotypic comparisons revealed that the different culture conditions and maturation stages did not impact the overall proliferation, differentiation, migration, and survival of hiPSC-NSCs *in vitro* and upon intracerebral transplantation in neonatal and adult immunodeficient mice^34^. Indeed, we reported similar engraftment and rostro-caudal migration of hiPSC-NSCs and hfNSCs upon transplantation in a mouse model of metachromatic leukodystrophy (MLD)^34^, a severe genetic demyelinating disease. The present study supports these findings, showing stable and robust engraftment and survival of hiPSC-NSCs up to 10 months after transplant into neonatal immunodeficient wild-type (WT) mice. The enrichment in NESTIN^+^ human cells detected within or close to the murine SVZ neurogenic niche suggests the preferential engraftment of hiPSC-NSCs in a microenvironment that might favor their proliferation, maturation, and long-term survival. Engrafted hiPSC-NSCs widely disperse and migrate in different CNS regions, with a preferential rostral migration. In contrast, transplantation in MLD pups led to a prominent caudal distribution of engrafted cells, possibly driven by neurodegeneration (e.g., of the cerebellum) and sulfatide accumulation (e.g., in white matter regions) associated with the pathology^34,163^. Engrafted hiPSC-NSCs predominantly differentiate into SOX10^+^ glia progenitors, S100β^+^ astrocytes, and GSTπ^+^ oligodendrocytes, with a minor commitment toward the neuronal lineage. We speculated that the glial skewing is not dependent on the pathological environment but is instead mostly related to the cell type composition/identity of the hiPSC-NSC population generated with this protocol, being displayed by hiPSC-NSCs engrafted in both WT (this study) and MLD^34^ mice. Of note, hiPSC-derived NE showed a propensity to differentiate into neurons upon transplantation in immunodeficient MLD^33^. Further experiments are required to clarify whether the preferential generation of glial cells described in our study is associated with increased proliferation/survival of MRG/LRG/glial progenitors in the donor hiPSC-NSC population and/or with *in vivo* maturation of ERGs. Interestingly, transplantation of hiPSC-derived rosette-type primitive neural progenitors into neonatal mice lead to an initial generation of neuroblasts at 3 weeks post-transplant, followed by the generation of astrocytes and oligodendrocyte progenitors at 6-13 months, suggesting that engrafted cells could acquire a gliogenic potential over time^164^.

One of the major concerns in the clinical translation of hiPSC-derived products for cell therapy is the safety risk associated with reactivation of pluripotency and cancerogenic genes^127^ or the activation of transcriptional and epigenetic programs leading to aberrant differentiation^39^. Our data show that hiPSC-to-NSC differentiation generated cells with distinct transcriptional and epigenetic profiles as compared to parental hiPSCs, characterized by a robust downregulation of pluripotency genes and factors driving mesodermal/endodermal specification and the loss of enhancers and SEs enriched in predicted binding sites of master regulators of pluripotency (OCT4, NANOG, c-MYC). Of note, scRNA-seq analysis confirmed the absence of iPSC contaminants expressing the pluripotency master regulator OCT4 or hiPSC/ESC markers (VRTN, ZSCAN10, LINC00678, and ESRG)^41^ in the three hiPSC-NSC clones analyzed, although residual expression of some pluripotent genes (e.g. CNMD1, DPPA4 and L1TD1) was detected at low levels. We report here the downregulation of pathways positively regulating the cell cycle and related processes (DNA replication processes and telomere extension) and driving cancer formation at both transcriptional and epigenetic levels. We observed the silencing of enhancers involved in MYC-associated pathways, which are a central hub in pluripotency maintenance^165^ and cancer formation^149^. The transcriptional and epigenetic switch-off of the MYC-pathway during hiPSC neural commitment is associated with an upregulation of the MYC-repressor MXD1^166,167^ soon after neural induction, further confirming the silencing of the self-sustained MYC-associated program. Global transcriptional analysis of hiPSC-NSCs, hfNSCs, and GSCs showed a low similarity in the transcriptional profiles between the neural populations and the aggressive glioblastoma cells^132^. The majority of potential oncogenic TFs, signaling molecules, and metabolic processes driving glioblastoma reprogramming and growth were expressed at significantly lower levels in hiPSC-NSCs compared to GSCs; instead, expression levels were similar to hfNSCs, which are characterized by absence of tumorigenic potential established in several clinical trials^25–28^. These *in vitro* observations on the safety of transplantable hiPSC-NSCs were corroborated by long-term *in vivo* studies demonstrating the absence of hyperproliferation or tumor formation in 10-month mice transplanted as neonates. Of note, the percentage of Ki67^+^ proliferating cells remained low and stable over-time, as compared to data collected in short-term (3- and 6-months) studies^34^.

Overall, our comprehensive omics analyses define hiPSC-NSCs as a heterogenous RG population composed of cells in a transient stage of maturation between MRG and LRG. Despite observing clone-related differences in overall cell composition, we did not detect OCT4^+^ hiPSC contaminants in any of the samples analyzed, which displayed instead the acquisition of a peculiar NSC signature similar to that shown by somatic hfNSCs (e.g., expression of key genes and pathways regulating growth factor-dependent proliferation and survival). The long-term transplantation study demonstrates the robust and stable engraftment of hiPSC-NSCs and their preferential gliogenic potential without major safety concerns associated with hyperproliferation or reactivation of a pluripotency program. The combination of our RNA-seq, ChIP-seq, and scRNA-seq analyses with data collected in ESC-derived neural cells could provide reference datasets to establish the heterogeneity, maturation stage, and cell identity of hiPSC-derived neural cells generated by applying different protocols and define the expression threshold of potential oncogenic genes in perspective of their clinical translation in cell therapies for neurodegenerative disorders.

## MATERIALS AND METHODS

### Isolation and culture propagation of hNSC lines

Human cells were used according to the guidelines on human research issued by the ethics committee of Ospedale San Raffaele, in the context of the protocol TIGET-HPCT.

The hiPSC clones were obtained from fibroblasts from the Cell Line and DNA Bank of Patients affected by Genetic Diseases (Institute Gaslini, Genova, Italy, http://www.gaslini.org) for HD1 (adult) or acquired from Invitrogen (C0045C/ Invitrogen) for HD2 (newborn) and characterized as previously described^34^.

hiPSC-NSC were generated from hiPSCs as previously described^34^. Briefly, hiPSCs were detached with dispase (Thermo Fisher Scientific) and cultured as embryoid bodies (EBs) in EB medium. On day 4, EBs were plated on Matrigel (BD Biosciences)-coated dishes and grown in EB medium supplemented with NOGGIN (250 ng/mL, R&D Systems). At day 10, medium was replaced with EB medium supplemented with Sonic Hedgehog (SHH; 20 ng/mL, R&D Systems) and fibroblast growth factor 8 (FGF8; 100 ng/mL, R&D Systems). Upon appearance of rosette-like structures (day 14), medium was changed to BASF medium (brain-derived neurotrophic factor [BDNF], ascorbic acid, SHH, and FGF8). On day 22, FGF8 was withdrawn, and cells were maintained in BAS medium (BDNF, ascorbic acid, and SHH). At day 29, cells were detached with Accutase (Thermo Fisher Scientific) and plated on poly-L-ornithine (20 μg/mL, Sigma-Aldrich)/laminin (10 μg/mL, Thermo Fisher Scientific)-coated dishes in hiPSC-NSC proliferation medium^34^.

The hfNSC line used here was originally isolated from fetal brain tissue (diencephalon/telencephalon; 10.5- week gestational age) obtained from Advanced Bioscience Resources, Inc., Alameda, CA, USA, and cultured in mitogen-supplemented serum-free medium as described^40^. For RNA-seq and ChIP-seq analyses, we used 3 replicates consisting of cells harvested at different subculturing passages (p19, p23, p25).

### Differentiation of hiPSC-NSCs in mixed populations of glial and neuronal cells

To obtain a mixed neuronal/astroglial population for SREBF1 KO experiments, we applied a previously described protocol^34^. Briefly, hiPSC-NSCs were detached as single cells and plated on Matrigel-coated dishes in mitogen-supplemented Neural Differentiation Medium^34^. After 3 days, cells were detached using Accutase and plated at a density of 20,000 cells/cm^2^ in Neural Differentiation Medium on Matrigel-coated dishes or Matrigel-coated coverslips and cultured for 15 days.

### Generation of SREBF1 knockout hiPSC clones

hiPSCs (clone HD1.3) were manually picked and plated on Matrigel-coated dishes in hiPSC medium supplemented with 10 µM Y27632 (Sigma) and kept in culture for 2 passages. When iPSCs reached 70% confluence, cells were detached with Accutase. Cas9 2NLS protein was incubated for 10 min at room temperature (RT) with a mixture of 3 synthetic gRNAs targeting SREBF1 exon 5 (CRISPR Gene Knockout Kit v2, Synthego) to form ribonucleoprotein (RNP) complexes (gRNA are listed in Suppl. Table 5). 2 × 10^5^ cells were resuspended in 17 μL of P3-supplemented nucleofection buffer (Lonza). We then added RNP mix, and cells were immediately nucleofected with Amaxa 4D-Nucleofector (Lonza), using program CB-150. After recovery (5-10 min at 37°C), nucleofected cells were collected and plated on a 15-cm mitotically inactivated murine embryonic fibroblast (MEF)-coated plate in hiPSC medium supplemented with 10 µM Y27632, while a subset of hiPSCs were plated on Matrigel for KO efficiency analysis. After colony formation, ∼40 colonies were picked and plated, with each clone in 1 well of a 96-well plate. After 5 days, one half of each colony was plated in MEF-coated 48-well plates and the other half in Matrigel-coated 48MW. Matrigel colonies were used for KO efficiency analysis and MEF colonies for amplification and cryopreservation. KO efficiency analysis was performed using the Synthego ICE Analysis Tool. Briefly, we designed primers flanking gRNA sites (Suppl. Table 5), PCR-amplified target regions from each clone, sequenced with the same primers used for PCR and performed analysis. Selected clones have KO-score = 100.

### Bulk RNA sequencing (RNA-seq) and data analysis

Total RNA samples were extracted using miRNeasy Mini Kit (Qiagen) according to manufacturer instructions and checked for integrity by microcapillary electrophoresis on a 2200 TapeStation instrument (Agilent Technologies). RNA-seq libraries were prepared using the Illumina TruSeq Stranded mRNA Library Prep Kit (Illumina) according to manufacturer instructions and sequenced on a HiSeq 2500 platform (Illumina) in a 125-cycle paired-end run at IGA Technology Services Srl (Udine, Italy). After performing fastq quality control by using the FastQC tool, raw reads were mapped to the human reference genome (hg19) using STAR aligner (v2.5.0a), and gene counts were calculated by HTSeq (v.0.6.1), using the hg19 Encode-Gencode GTF file (v19) as gene annotation file. Raw read counts were used as input for global data visualization and differential expression analysis by DESeq2 Bioconductor package (v1.8.1) in R environment (v3.2.2). Raw counts were normalized using DESeq2 rlog function and then used to perform sample-to-sample distance visualizations (using the DESeq2 plotPCA, sampleDists, and pheatmap functions). For differential gene expression comparison of hiPSC-NSC vs. hiPSC and hiPSC-NSC vs. hfNSC, |log2FC| > 1 and adjusted *p*-value (padj, Benjamini-Hochberg correction) < 0.05 were used as the cut-off to define statistically significant differentially expressed genes (DEGs). For comparison with gene expression profiles of neural samples by Ziller et al.^67^ and glioblastoma samples by Park et al.^132^, fastq files were downloaded from the Gene Expression Omnibus repository (GEO Accession Code GSE62193 for Ziller et al^67^.; GEO Accession Codes GSE87617 and GSE87615 for Park et al.^132^), and then mapped, counted, and normalized together with our samples as described above.

### Chromatin immunoprecipitation

hiPSCs, hiPSC-NSCs, and hfNSCs were crosslinked for 10 min at RT with 1% methanol-free formaldehyde-containing medium (Fisher Scientific, Leicestershire, UK) and blocked with 0.125 M glycine. Crosslinked cells were washed with PBS containing protease and phosphatase inhibitors and stored at -80°C until use. Nuclei were collected upon cell lysis using Cell Lysis Buffer and nuclei release via dounce homogenization with a tight pestle. Nuclear extracts were sonicated in 1.5 mL Diagenode tubes on a Bioruptor Pico in Sonication Buffer for 12 (for hiPSCs) or 15 (for hiPSC-NSCs and hfNSCs) cycles of 30 sec ON / 30 sec OFF to obtain DNA fragments averaging 200 bp in length (checked with microcapillary electrophoresis on a 2200 TapeStation instrument). Chromatin preclearing was performed using IgG isotype (1 ug IgG/10^6^ cells)-Protein G Agarose-coated beads (Thermo Fisher Scientific) mixed with chromatin. The equivalent of 3-6 x10^7^ cells were precipitated overnight with 1 ug IgG/10^6^ cells of rabbit antibody against H3K27ac (ab4729, Abcam). Before antibody precipitation, total chromatin was collected as input. H3K27ac^+^ chromatin was precipitated with precleared (RIPA Buffer + protease inhibitors; BSA at 0.5 ug/ul and S256 containing salmon sperm at 2.7 ug/ul overnight at 4°C) Protein G Agarose-coated beads for 2 hours on a rotating wheel at 4°C. Beads were then washed with RIPA buffer, LiCl buffer, and TE buffer. Chromatin was eluted with Elution Buffer, and reverse crosslinking was performed with 0.3 M NaCl and RNase A (10 mg/mL) for 4-5 hrs at 67°C. Chromatin was precipitated overnight in 100% ethanol with glycogen (5 mg/mL) at -20 °C. After protein digestion (PK Buffer and Proteinase K (20 mg/mL) for 2 hrs at 45°C) DNA was purified with QIAquick PCR Purification Kit (Qiagen, Germany) according to manufacturer instruction. Real-time SYBR Green PCR was used to validate genomic regions enriched in H3K27ac in each cell type. Primers were derived from Rada Iglesias *et al.*^151^ (Suppl. Table 6). A negative control (Neg; genomic region that falls in a “gene desert” region), a double-positive control (FGFR1, acetylated in both hiPSCs and hiPSC-NSCs), a positive control for hiPSCs (Lin28), and a positive control for hiPSC-NSCs (PPAP2b) were chosen.

### ChIP-seq library preparation, sequencing, and analysis

Illumina libraries were prepared from 10 ng of immunoprecipitated DNA (IP) and control DNA (INPUT: nuclear extracts sonicated but not immunoprecipitated) following the Illumina ChIP-Seq DNA Sample Prep Kit. Libraries were checked by capillary electrophoresis on an Agilent 2100 Bioanalyzer with the High Sensitivity DNA assay and quantified with Quant-iT PicoGreen dsDNA Kits (Invitrogen) on a NanoDrop Fluorometer. Each library was sequenced in one lane of a single-strand 50 bp Illumina run. Raw reads were mapped against the human reference genome (build hg19) using Bowtie^168^, allowing up to 2 mismatches (-v 2 option) and discarding multiple alignments (-m 1 option). Each BAM file was then processed by using SAMtools^169^ and converted into a bed file using BEDTools^170^. The quality of each sequenced sample was checked using cross-correlation analysis implemented in the spp R package^171^. ChIP-seq peak calling was performed with MACS2 (with –broad and --qvalue 0.05 options)^172^ and using each INPUT data to model the background noise. A custom R-workflow was developed to identify promoters and enhancers. The pipeline analyses merged H3K27ac^+^ broad peaks generated by MACS2 on a cell basis and then identified putative promoters (< 2kb from transcription start site [TSS]) and enhancers (> 2kb from TSS) if present in at least 2 replicates for each condition. Promoters were annotated to the RefSeq gene whose TSS was the nearest to the center of the H3K27ac peak. For enhancers, windows of 100, 200, and 400 kb from enhancer boundaries were designed, and the RefSeq genes falling into these windows were identified. When comparing two conditions, promoters or enhancers were defined as specific if they did not overlap up to a fraction of 0.3, reciprocally. Enhancers were stitched and Super Enhancers were defined using ROSE code (https://bitbucket.org/young_computation/rose), as previously described in^173,174^. Briefly, this algorithm stitches enhancers together if they lie within a certain distance and ranks the enhancers by their input-subtracted H3K27ac signal. It then separates Super Enhancers from typical enhancers by identifying an inflection point of H3K27ac signal vs. enhancers rank. ROSE was run with a stitching distance of 12,500 bp. In addition, all the enhancers wholly contained in a window ± 2,500 bp around an annotated TSS (RefSeq, build hg19) were excluded from stitching, allowing for a total 5,000 bp promoter exclusion zone. For Super Enhancers, windows of 100, 200, and 400 kb from enhancer boundaries were designed, and the RefSeq genes falling into these windows were identified. Specific Super Enhancers were defined in the same way as specific promoters and enhancers. TF motif finding was performed using HOMER^175^. Background sequences were automatically selected and weighted to resemble the same GC-content distribution observed in the target sequences. Top enriched motifs were shown. Motifs of TFs not expressed in these cell types, enriched in < 5% of the target sequences, or associated with a *p*-value > 10^−2^ were excluded.

### Gene Ontology analysis of bulk RNA-seq and ChIP-seq datasets

For RNA-sequencing, we performed gene ontology (GO) analyses considering a log2 Fold change of > 1 or < -1 with an adjusted *p*-value < 0.05 for each comparison (hiPSC-NSC vs. hiPSC, hiPSC-NSC vs. hfNSC, hiPSC-NSC vs. GSC, and hfNSC vs. GSC). We used ToppFun (ToppGene suite)^176^ to identify enriched biological processes or pathways, using the Bonferroni correction with 0.05 as the significance cut-off level. We represented the most significant and/or relevant pathways or biological processes ranked on the highest Bonferroni Q value (when applicable) or *p*-value. For Ingenuity Pathway Analysis (Qiagen, Germany), we used the dataset of DEGs in the comparison hiPSC-NSC vs. hiPSC with a log2 Fold change of > 1.5 or < -1.5 and an adjusted *p*-value < 0.01. We then selected upstream regulators with a predicted activation state concordant with gene expression and with a *z*-score > 2 or < -2 for activated or inactivated TFs, respectively. For enhancer and Super Enhancer GO analyses, we considered a window of 400 kb from the boundaries (supervised analysis). We then mapped the RefSeq Genes contained in the windows and identified these as enhancer- or Super Enhancer-related genes, picking only the genes whose expression was concordant with the cell-specificity of the regulatory regions (i.e., for hiPSC enhancers, we picked genes contained in the 400 kb windows whose expression had a fold change < 1.5 with an adjusted *p*-value < 0.05). We then ran GO analysis via ToppFun as described above.

### Single cell-RNA sequencing

For single cell-RNA seq experiment hiPSC-NSC were enzymatically dissociated with Accutase while hfNSC were mechanically dissociated by pipetting. Cells were then checked for viability (>90%) counted and resuspended at a concentration of 1,000 cells/μl.

Single cells were processed using the Chromium Single Cell 3’ Reagent Kit (v3.1 Chemistry) and the 10x Chromium Controller platform (10X Genomics). The target cell recovery was 2000 cells. Samples were processed following the manufacturer’s protocol. The resulting libraries were assessed for size distribution and concentration using the HS DNA assay (Agilent) on the TapeStation platform (Agilent). Libraries were sequenced on a Novaseq 6000 (Illumina) platform aiming at 50’000 reads per single cell.

CellRanger v6.1.1 software (10X Genomics) was used to perform demultiplexing of the input files, alignment to the human reference genome (GRCh38) using the STAR software, and UMI quantification to produce a cell-by-gene matrix for all the samples, which were then imported in the R environment (v.4.1.3) and analysed with Seurat (v4.1.1). In details, cells expressing less than 200 (indication of low-viability) or more than 6000 (indication of doublets) genes, as well as those having more than 20% of transcripts coming from mitochondrial genes (indication of dying cells), were removed. Samples were then merged, and the resulting combined object was analyzed by performing log-normalization with a scale factor of 1000 by using the NormalizedData function of the Seurat package, principal component analysis, batch removal (using Harmony), clustering, and Uniform Manifold Approximation and Projection (UMAP) embedding computation. Cluster markers were computed by using the FindAllMarkers function of the Seurat package, which exploits a Wilcoxon Rank Sum test for significance. The Seurat function AddModuleScore was used to test signatures coming from different datasets^37,61,67,106–108^ by computing the average expression in different set of cells (i.e., samples or clusters), while the R/Bioconductor package clusterProfiler (v4.7.1) was employed to perform enrichment of cluster markers on the KEGG database. Single-cell pseudotime trajectories were computed with the R package Slingshot (v2.2.1) which exploits previously computed cell clusters, and focusing on the transcriptional lineage that describes the progressive transition from cluster 1 to cluster 7 by also identifying genes whose expression changes along that trajectory.

### Gene expression analysis by qRT-PCR

Total RNA was extracted from cells using the RNeasy Mini Kit (Qiagen) according to the manufacturer instructions and retrotranscribed with the QuantiTect Reverse Transcription Kit (Qiagen). Quantitative real-time PCR reactions (TaqMan and SYBR Green) were performed on a ViiA 7 Real-Time PCR System (Applied Biosystems). For quantitative real-time PCR of selected TFs, we used customized TaqMan Array 96-Well Fast Plates (TaqMan Gene Expression Assays, Thermo Fisher) with pre-spotted lyophilized primers + probes mix. SYBR Green primers and TaqMan probes are listed in Suppl. Table 7.

### FACS analysis

hiPSC colonies were manually picked on Matrigel-coated dishes and expanded in hiPSC medium supplemented with 10 µM Y27632. hiPSCs and hiPSC-NSCs were detached with Accutase, whereas hfNSC neurospheres were mechanically disaggregated to single-cell suspensions by pipetting. The following antibodies were used: anti-NGFR APC-conjugated antibody (560326, BD Pharmingen), anti-SSEA4 APC-conjugated antibody (FAB1435a, R&D Systems). For Annexin V/7AAD analysis, we used Dead Cell Apoptosis Kits with Annexin V for Flow Cytometry (Thermo Fisher) according to manufacturer instructions. Unstained cells were used as negative control to define the gate strategy (SSC, FSC, laser intensity parameters). Raw data were analyzed with FlowJo software version 10.8.1.

### Mice

*C57BL/6; Rag^-/-^; γ-chain^-/-^* mice were maintained in the animal facility at the San Raffaele Scientific Institute, Milano, Italy. Mice were housed in microisolators under sterile conditions and supplied with autoclaved food and water. All experiments and procedures described in this study were performed according to protocols approved by the internal Institutional Animal Care and Use Committee and reported to the Italian Ministry of Health, as required by Italian law (IACUC # 931).

### Cell transplantation, tissue collection and processing

hiPSC-NSCs (clones HD 1.1, HD 1.3, HD 2.2) were dissociated to single cells with Accutase, washed with PBS, and resuspended in sterile-filtered PBS + 0.1% DNase I to a concentration of 100,000 cells per μL. Cells were transplanted into post-natal day (PND) 2-4 mice by bilateral intracerebroventricular injection (100,000 cells per 1 μL per injection site) using a 33G needle-Hamilton syringe as previously described^177^. At 10 months after transplantation, mice were euthanized after anesthetic agent overdose by intraperitoneal injection and intracardially perfused with 0.9% NaCl plus 50 U/mL heparin. Brain and spinal cord were collected and post-fixed in 4% paraformaldehyde in PBS for immunofluorescence analysis. Fixed tissues were cut at the vibratome to obtain sagittal and coronal free-floating sections (thickness 40 μm).

### Immunofluorescence and confocal analyses

Immunofluorescence on free-floating vibratome sections was performed as previously described^177^. Tissue sections were rinsed with PBS, incubated in blocking solution (10% NGS + 0.3% Triton X-100 in PBS) for 1 hr at RT and then with primary antibodies overnight at 4°C. Samples were then incubated for 2 hrs at RT with species-specific fluorophore-conjugated secondary antibodies in 1% NGS in PBS. Primary and secondary antibodies are listed in Suppl. Table 8. Nuclei were counterstained with Hoechst 33342 (Invitrogen). Sections were mounted on glass slides using FluorSave (Calbiochem). No detectable signal was observed in samples lacking primary antibodies. Confocal images were acquired at different magnifications with a TCS SP8 confocal microscope (Leica) or an RS-G4 upright confocal microscope (MAVIG Research). Data were analyzed with FIJI software (ImageJ, National Institutes of Health)^178^, LAS X software (Leica Application Suite X, RRID: SCR_013673), and Photoshop CS4 (Adobe).

### Evaluation of cell engraftment and phenotypic characterization of engrafted cells

Human cells were identified by using nuclear (anti-hNuclei) and cytoplasmic (STEM121) anti-human antibodies. hiPSC-NSC engraftment was estimated in coronal and sagittal brain sections (12-16 sections per mouse, corresponding to one out of six series) by using anti-hNuclei antibody. An automated count of nuclei was performed with FIJI on masks of confocal images acquired at 20x magnification using the RS-G4 upright confocal microscope on the whole section. The number of hNuclei^+^ cells per section was multiplied by 6 to obtain an estimate of the total number of engrafted cells per brain. The engraftment percentage was expressed as (total number of engrafted cells / total number of transplanted cells) × 100. The distribution of engrafted cells was assessed by automated counting of human cells in coronal telencephalon sections organized along the rostro-caudal axis. Engrafted cells were characterized by immunofluorescence followed by confocal microscopy analysis in brain sections (at least three fields per region per hemisphere) using anti-hNuclei or - STEM121 antibodies coupled with antibodies against lineage-specific markers and nuclear counterstaining. Z-stacks were recorded at 40X magnification using the TCS SP8 confocal microscope.

### Statistics

Data were analyzed with GraphPad Prism for Windows version 8.0a and expressed as the mean ± standard error of the mean (SEM). One-way ANOVA followed by appropriate post-tests and Mann-Whitney tests were used; *p*-value threshold for statistical significance was considered 0.05. Pairwise Wilcoxon test was used to determine significant differences in the expression values associated with enhancers and Super Enhancers in the different cell types. Adjusted *p*-value was used to determine statistical significance in DEGs analysis (padj < 0.05). Bonferroni Q value was used to determine statistical enrichment in ToppFun gene ontology analyses (q-val < 0.05). The number of samples and statistical tests used are indicated in the figure legends.

## Supporting information

Supplementary Informations

## Author contribution

A.M., V.M., and A.G. designed and conceived the study;

M.L. performed in vitro and in vivo experiments;

C.G. performed in vivo experiments;

S.B., I.M, L.P, I.C., C.P, M.L. and V.M analyzed the scRNA-seq data, RNA-seq and ChiP-seq data;

A.M. provided expertise and resources for conducting RNA-seq and ChiP-seq analysis;

A.G. and V.M. provided expertise, resources, and supervised the whole study;

V.M. and M.L. wrote the manuscript, with critical input and final approval of A.G.

## Acknowledgments

We thank Eleonora Ciccarelli and Oriana Romano for help with sample collection and processing; Cesare Covino for microscopy support; Francesca Giannese for RNAseq support; Stefano Pluchino, Rossella Galli, and Giorgia Quadrato for critical review of the paper; Daniel Ackerman (Insight Editing London) for professional editing of the manuscript. Part of this work was carried out in the core facilities established at IRCSS Ospedale San Raffaele and Università Vita-Salute San Raffaele: ALEMBIC (Advanced Light and Electron Microscopy BioImaging Center); COSR (Center for Omics Sciences); FRACTAL (Flow cytometry Resource, Advanced Cytometry Technical Applications Laboratory).

This study was funded by grants from Fondazione Telethon (TGT16D02; TTAGD0222TT) and Italian Ministry of Health, Ricerca Finalizzata (RF-2016-02362404) to A.G; French State funding from the Agence Nationale de la Recherche “Investissements d’Avenir” program (ANR-10-IAHU-01) to A.M. The sponsor(s) had no role in the study design or the collection, analysis, and interpretation of data; or the decision to submit the article for publication. M.L. conducted part of this study to fulfill his PhD in Molecular Medicine, XXXI cycle (Universita’ Vita-Salute San Raffaele) with the support of fellowships from Fondazione Telethon and Erasmus Plus Programme.

## Data Availability

Bulk RNA-seq and ChIP-seq data have been deposited at GEO under accession number GSE239446. Single cell RNA-seq data have been deposited at GEO under accession number GSE238206.

Analyzed RNA-seq, ChIP-seq and scRNA-seq data (list of DEGs and GO terms) (Suppl. Files 1-3) are available upon request.

