## Supplementary Informations for "Human iPSC-derived neural stem cells display a radial glia-like signature *in vitro* and favorable long-term safety in transplanted mice"

#### Supplementary information

- **Supplementary Figures**

Supplementary Figure 1. Expression of NSC markers and predicted TF network in hiPSC-NSCs.

Supplementary Figure 2. Expression of TFs during hiPSC neural differentiation and RG markers in hiPSC-NSCs.

Supplementary Figure 3. Validation of ChIP-seq analysis in hiPSCs, hiPSC-NSCs, and hfNSCs.

Supplementary Figure 4. Comparable activation of cell cycle and metabolic pathways in hiPSC-NSCs and hfNSCs.

Supplemental Figure 5. Single-cell RNA-seq analyses in hiPSC-NSCs and hfNSCs.

Supplementary Figure 6. Role of SREBF1 in the astroglial commitment of hiPSC-NSCs.

Supplementary Figure 7. Engrafted hiPSC-NSCs migrate along the rostro-caudal axis and primarily differentiate into glia progenitors.

- **Supplementary Tables**

Supplementary Table 1. Summary of hiPSC clones generated by reprogramming of healthy donor (HD) fibroblasts.

Supplementary Table 2. Upstream regulators identified by Ingenuity Pathway Analysis (IPA) on RNA-seq analysis of hiPSC-NPCs vs. hiPSCs.

Supplementary Table 3. Upstream regulators identified by Ingenuity Pathway Analysis (IPA) on the dataset of genes close to hiPSC- and hiPSC-NSC-specific enhancers

Supplementary Table 4. Upstream regulators identified by Ingenuity Pathway Analysis (IPA) on RNA-seq analysis of hiPSC-NSCs vs. hfNSCs.

Supplementary Table 5. gRNA for SREBF1 KO and primers used for PCR amplification and sequencing.

Supplementary Table 6. List of primers used for qRT-PCR on immunoprecipitated chromatin.

Supplementary Table 7. List of primers and probes used for SYBR Green and TaqMan qRT-PCR.

Supplementary Table 8. List of primary and secondary antibodies with antigen, host species, provider, product number, and working dilutions indicated.

- **Supplementary Files (available upon request)**

Supplementary File 1. Analyzed RNA-seq data (list of DEGs and GO terms)
Supplementary File 2. Analyzed ChIP-seq data (list of DEGs and GO terms)
Supplementary File 3. Analyzed scRNA-seq data (list of DEGs and GO terms)

### Supplementary figures

Supplementary Figure 1

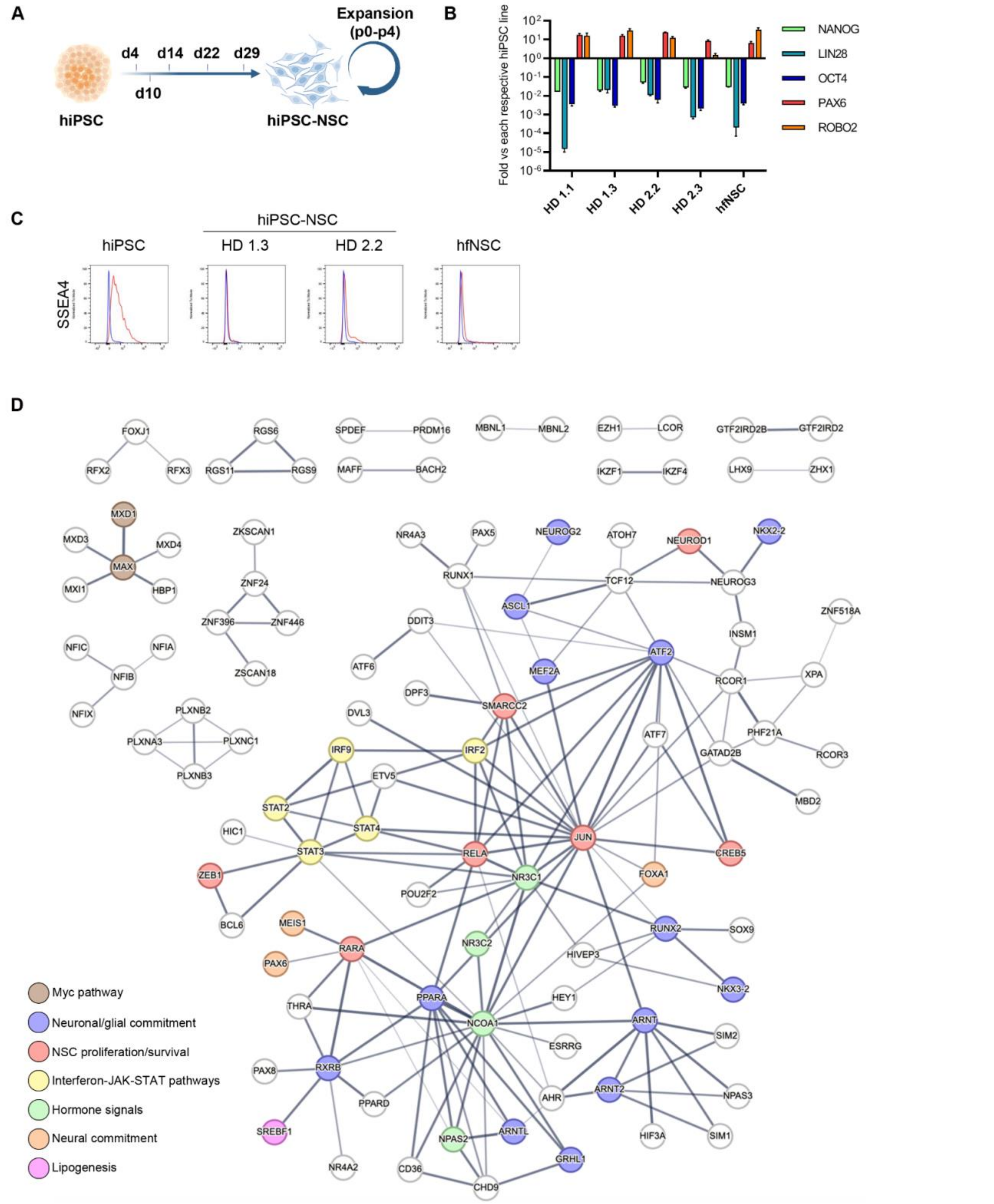


**Supplementary Figure 1. Expression of NSC markers and predicted TF network in hiPSC-NSCs. A)** Schematic representation of neural differentiation protocol. Timepoints analyzed in this study are shown and correspond to different stages during hiPSC-to-NSC commitment (hiPSCs, day 0; Embryoid bodies, day 4; early and late rosette-like formations, days 10 and 14; hiPSC-NSC maturation, days 22 and 29) and hiPSC-NSC expansion in growth media (passages 0-4). Created with BioRender.com. **B)** Bar plot showing the upregulation of NSC markers (PAX6, ROBO2) and downregulation of hiPSC markers (NANOG, LIN28, OCT4) in hiPSC-NSCs at levels similar to hfNSCs. For each gene, the expression levels in hiPSC-NSCs are reported as fold changes vs. the corresponding parental hiPSC clone, whereas expression levels in hfNSCs are reported as fold changes vs. the mean values in hiPSC clones. Data are expressed as mean ± SEM of n = 2-3 independent experiments. **C)** Representative FACS plots of SSEA4 (pluripotency marker) expression in hiPSCs, hiPSC-NSCs (clones HD 1.3 and HD 2.2), and hfNSCs. Blue lines, unstained cells; red lines, stained cells. **D)** Protein-protein interaction network functional enrichment analysis (STRING) of TFs upregulated in hiPSC-NSCs vs. hiPSCs ($log_2$ fold change ± 1, adjusted *p*-value < 0.05). Proteins are colored according to NSC functions defined based on published data, as described in Results.

**Supplementary Figure 2**

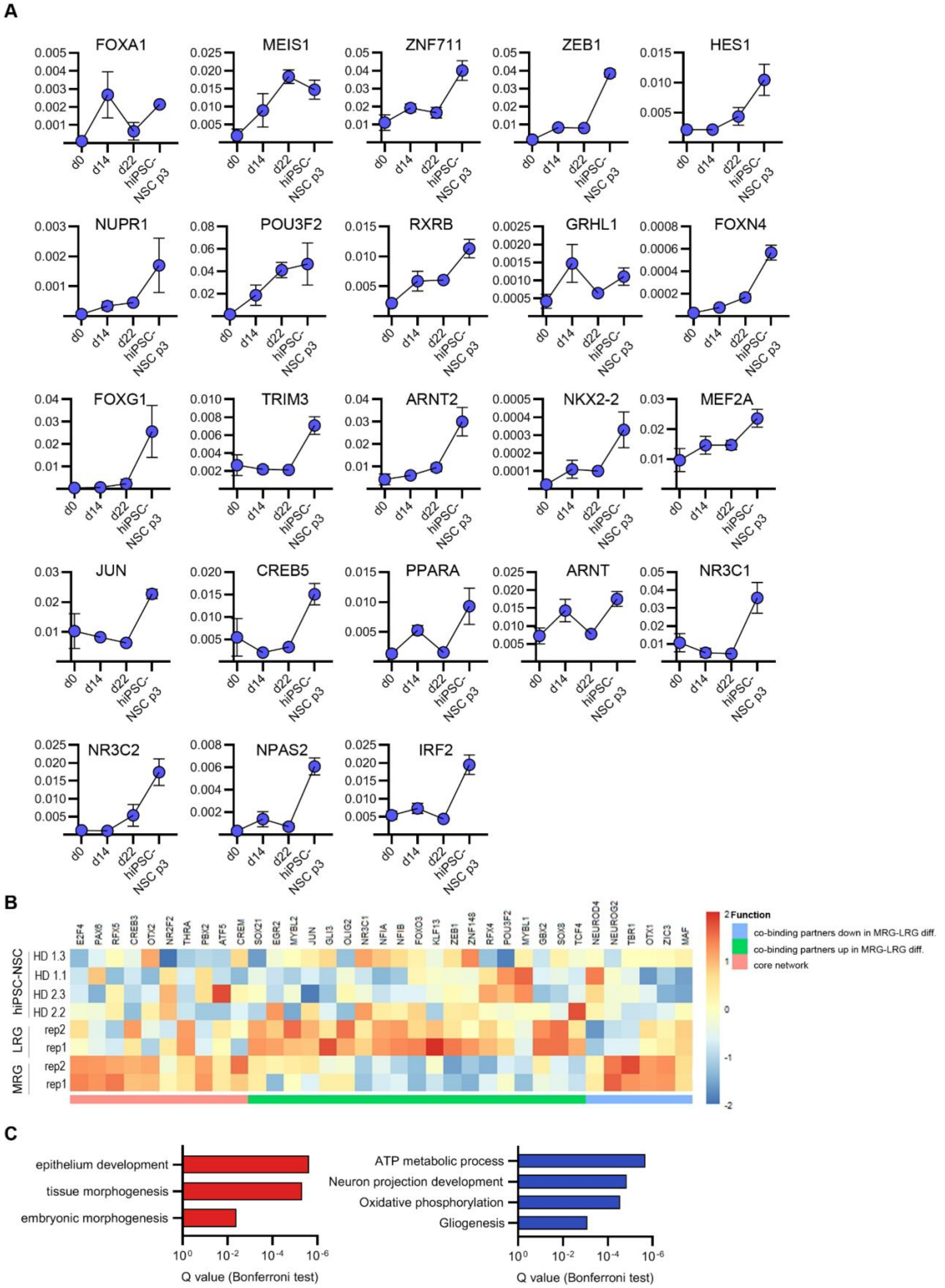


**Supplementary Figure 2. Expression of TFs during hiPSC neural differentiation and RG markers in hiPSC-NSCs. A)** Time-course qRT-PCR analysis showing the expression of TFs during hiPSC-to-NSC differentiation. Timepoints analyzed: hiPSCs (day 0), rosette-like formations (day 14), hiPSC-NSC maturation (day 22), and hiPSC-NSCs at passage 3. Expression levels are normalized on the housekeeping gene GAPDH. Each dot represents the mean ± SEM of 2 biological replicates. **B)** Heatmap showing the expression levels in hiPSC-NSCs of core and co-binding TFs regulating ESC neural commitment[67] in comparison to ESC-derived MRG and LRG. Color scale indicates the average expression levels of these genes in each cell population (blue, low; red, high). **C)** Bar plots show selected biological processes by gene ontology enrichment analysis of upregulated (red bars) and downregulated (blue bars) genes in hiPSC-NSCs vs. ESC-derived LRG[67] ($\log_2$ fold change ± 1, adjusted $p$-value < 0.05).

Supplementary Figure 3

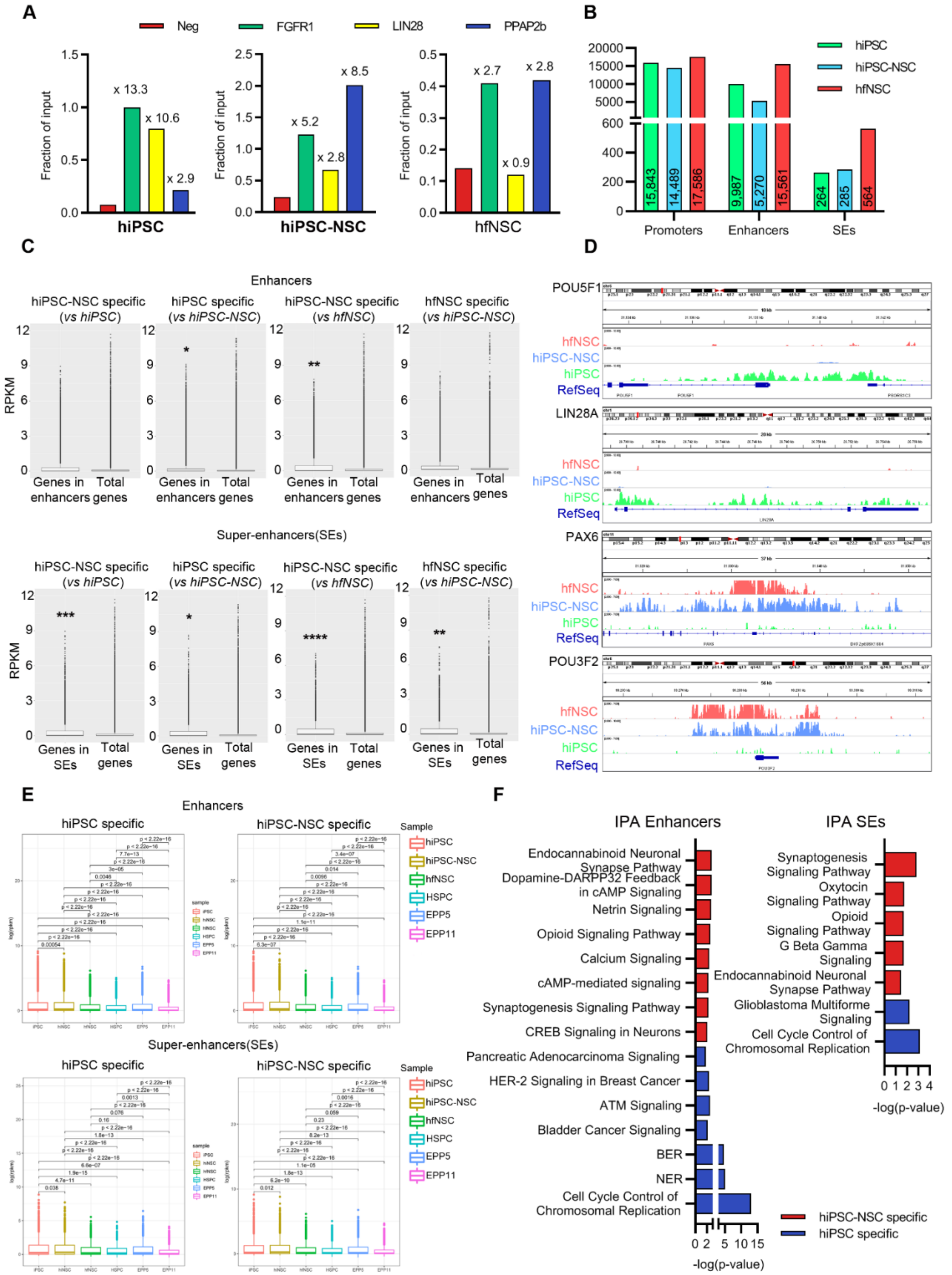


**Supplementary Figure 3. Validation of ChIP-seq analysis in hiPSCs, hiPSC-NSCs, and hfNSCs. A)** qRT-PCR on H3K27ac$^{+}$ immunoprecipitated chromatin for genomic regions containing pluripotent (Lin28) and NSC (PPAP2b) markers. FGFR1 serves as positive control in all cell populations while chr13:65364840+65364921 (82 bp) serves as negative control (Neg). Data are represented as fold change vs. the corresponding input sample (fraction of input). Enrichment on negative fraction (background) is indicated. **B)** Bar plot showing the number of $H3K27ac^+$ reads corresponding to promoters, enhancers, and SEs identified in each cell population. In each bar are indicated the numbers of $H3K27Ac^+$ reads. **C)** Box plot of expression levels (Reads Per Kilobase Million; RPKM) of genes close to cell-specific enhancers and SEs in comparison with total gene expression levels in each cell population. Welch Two-sample t-test, *, *p-adj* < 0.05; **, *p-adj* < 0.01; ***, *p* < 0.001, ****, p-adj < 0.0001 **D)** Integrative Genomic Viewer **(**IGV) snapshot of $H3K27ac^+$ peaks at pluripotency (POU5F1, LIN28A) and NSC (PAX6, POU3F2) genes in hiPSCs, hiPSC-NSCs, and hfNSCs. Genomic scale and RefSeq gene are indicated. **E)** Box plot of expression levels (logRPKM) of genes close to hiPSC- and hiPSC-NSC-specific enhancers and SEs detected by comparing hiPSC and hiPSC-NSC datasets. To verify the specificity of selected regulatory regions, the expression levels of selected genes were evaluated in published RNA-seq datasets retrieved from human stem/progenitor cells (HSPC) and erythroid progenitor/precursor cells at different stages of maturation (EPP5: day 5, EPP11: day 11)[80]. Pairwise Wilcoxon test was used to determine significant differences in the expression values between different cell types. **F)** Bar plots showing the IPA analysis of up- and downregulated genes ($\log_2$ fold change ± 1, adjusted *p*-value < 0.05) close to cell-specific enhancers (left plots) and SEs (right plots) in hiPSCs (blue bars) or hiPSC-NSCs (red bars).

**Supplementary Figure 4**

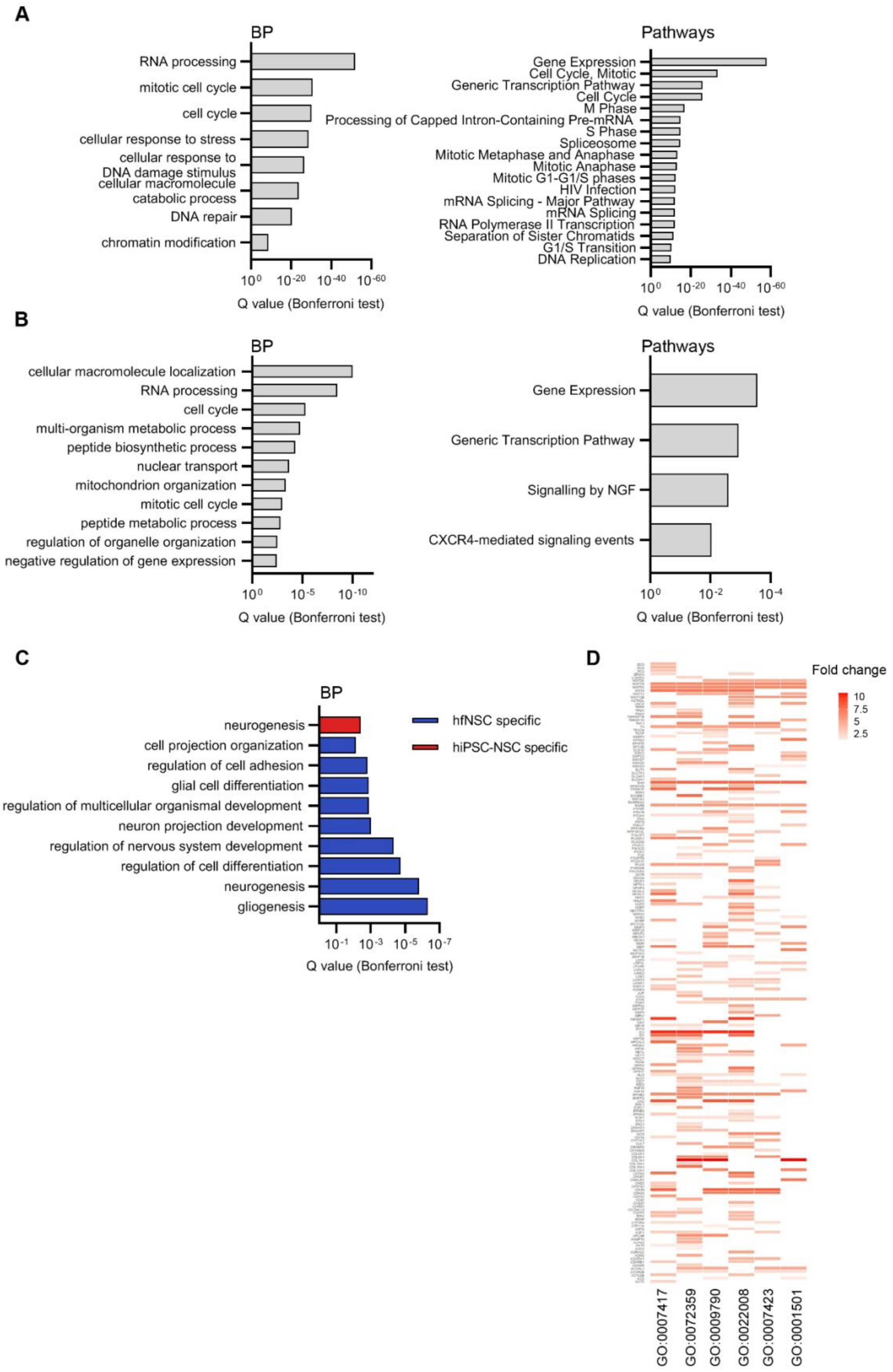


**Supplementary Figure 4. Comparable activation of cell cycle and metabolic pathways in hiPSC-NSCs and hfNSCs. A)** Gene ontology enrichment analysis of genes not differentially expressed between hiPSC-NSCs and hfNSCs detected in RNA-seq analyses. Bar plots indicate selected biological processes (BP, left plots) and pathways (right plots) similarly activated in the two neural populations. **B)** Gene ontology enrichment analysis of genes close to common enhancers in hiPSC-NSCs and hfNSCs. Bar plots show selected BP (left plots) and pathways (right plots) similarly activated in the two neural populations. **C)** Bar plot showing BP associated with genes close to hiPSC-NSC-specific (red bar) and hfNSC-specific (blue bars) SEs. **D)** Heatmap of Cluster Profiler analysis showing genes shared among GO terms associated with neural (GO:0007417, GO:0022008) and non-neural (GO:0072359, GO:0009790, GO:0007423, GO:0001501) biological processes. Fold change enrichment is indicated by the color scale (white, low; red, high).

#### Supplementary Figure 5

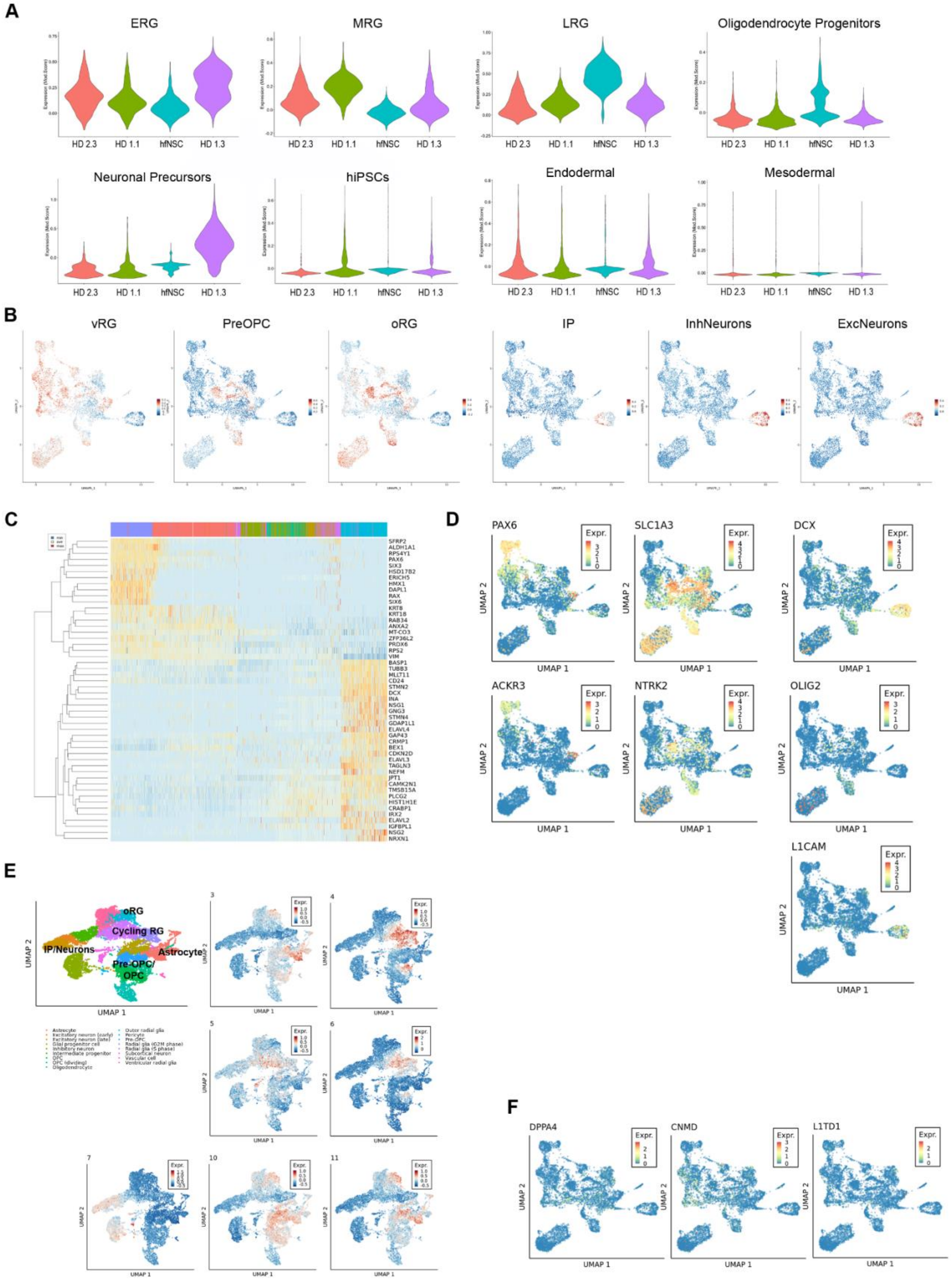


**Supplemental Figure 5. Single-cell RNA-seq analyses in hiPSC-NSCs and hfNSCs. A)** Violin plots showing the expression levels in hiPSC-NSCs (clones HD 1.1, HD 1.3, and HD 2.3) and hfNSCs of gene signatures associated with ERG, MRG, LRG, Oligodendrocyte Progenitors, Neuronal Progenitors, hiPSCs,

Endodermal cells, and Mesodermal cells. **B)** UMAP plot showing the distribution in scRNA-seq samples of cells expressing markers of cell populations isolated from fetal human brain (ventricular RG, vRG; outer RG, oRG; Pre-OPC; Intermediate Progenitors, IP; inhibitory neurons, InhNeurons; excitatory neurons, ExcNeurons)[108]. **C)** Heatmap indicating the variations in the expression levels of the 50 top genes along the pseudotime trajectory from Cluster 1 to Cluster 7. Expression is depicted according to the color scale (blue, low; red, high). **D)** UMAP plots showing the distribution of cells expressing membrane-bound markers (ACKR3, NTRK2, L1CAM in $PAX6^{+}$ early RG, $Slc1a3^{+}$ mature RG, $OLIG2^{+}$ OPC, and $DCX^{+}$ neuronal progenitors. **E)** UMAP plot of top 50 genes that identify hiPSC-NSC/hfNSC-derived mature RG (Clusters 3-6 and 11) and committed progenitors (clusters 7, 10) in scRNA-seq datasets of human fetal brain tissues[108]. **F)** UMAP plot showing minimal expression of hiPSC-associated markers (CNMD, DPPA4, and L1TD1) in scRNA-seq samples. In C and D color scale indicates the expression levels of each gene (blue, low; red, high).

**Supplementary Figure 6**

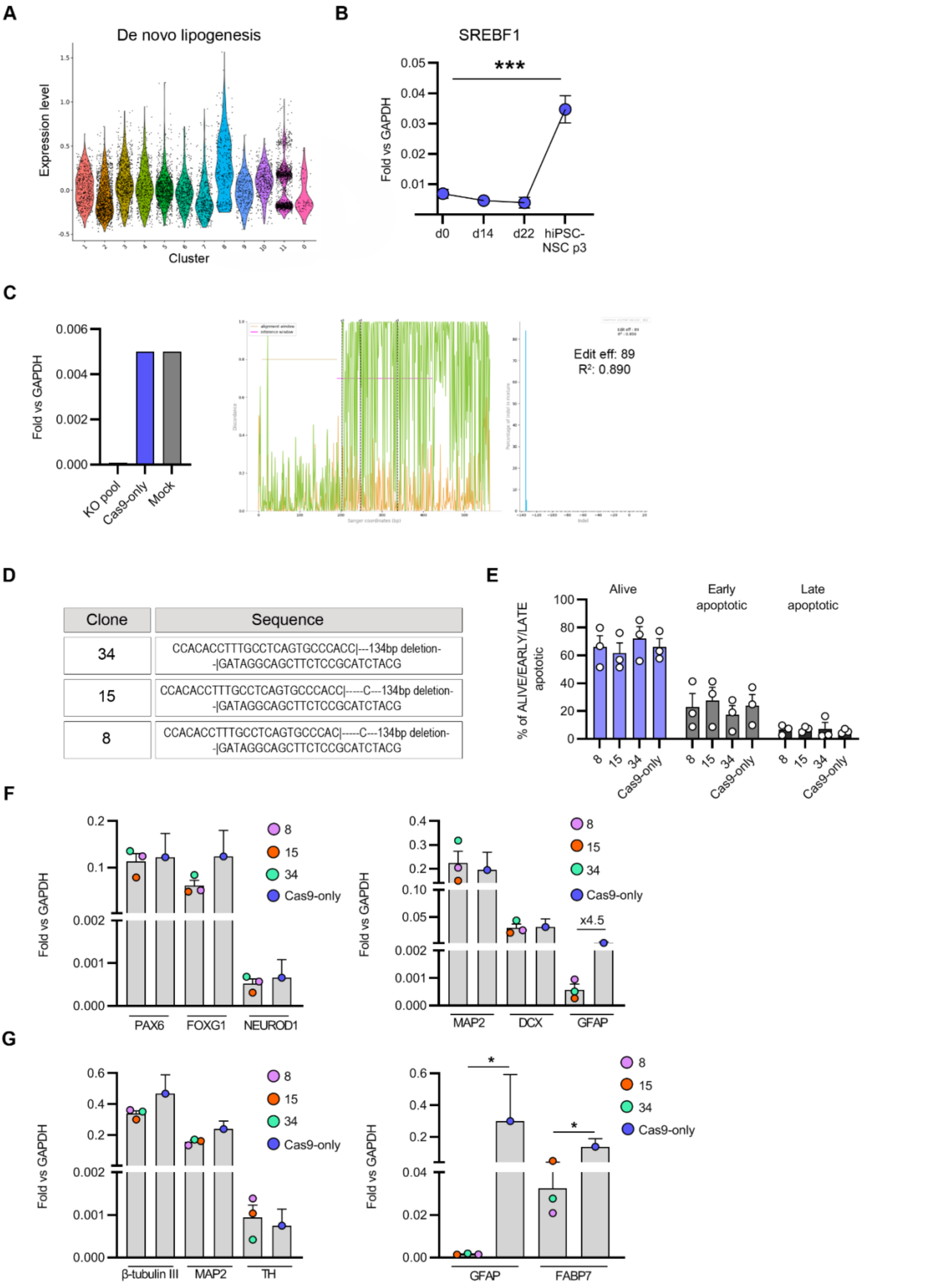


**Supplementary Figure 6. Role of SREBF1 in the astroglial commitment of hiPSC-NSCs. A)** Seurat violin plots showing the expression levels in the identified clusters of gene signature genes associated with *de novo* lipogenesis. **B)** Time-course qRT-PCR analysis of SREBF1 expression during hiPSC-to-NSC differentiation.

Timepoints analyzed: hiPSCs (day 0), rosette-like formations (day 14), hiPSC-NSC maturation (day 22), and hiPSC-NSCs at passage 3. Expression levels are normalized on the housekeeping gene GAPDH. Each dot represents the mean ± SEM of 3 biological replicates. One-way ANOVA followed by Dunn's multiple comparison test: ***, $p < 0.001$. **C)** Left: bar plot (left panel) showing the expression levels of SREBF1 mRNA in pooled SREBF1 KO hiPSCs as compared to control cells treated only with Cas9 protein (Cas9 only) and untreated samples (Mock). Expression levels are normalized on the housekeeping gene GAPDH. Right: results of ICE (Inference of CRISPR Edits) analysis showing the discordance plot of sgRNA target sequence and percentage of InDels on pooled SREBF1 KO hiPSCs. **D)** Table reporting the sequences of SREBF1 alleles in selected CRISPR/Cas9-edited hiPSC clones (clones 8, 15, 34). **E)** Annexin V/7-AAD FACS analysis to evaluate the percentage of living (Annexin $V^-$/7-$AAD^-$), early apoptotic (Annexin $V^+$/7-$AAD^-$), and late apoptotic (Annexin $V^+$/7-$AAD^+$) cells in SREBF1 KO hiPSC-NSC clones in comparison with Cas9-only treated cells. Data are expressed as mean ± SEM. Each dot represents a technical replicate. **F)** qRT-PCR analysis of the expression levels of NSC (PAX6, NEUROD1, FOXG1), neuronal (MAP2, DCX), and astrocytic (GFAP) markers in SREBF1 KO hiPSC-NSC clones and control cells (Cas9-only). Expression levels are normalized on the housekeeping gene GAPDH. Each dot represents the mean ± SEM of 2/3 technical replicates/clone. **G)** qRT-PCR analysis of the expression levels of pan-neuronal (β-tubulin III, MAP2), dopaminergic (TH), and astrocytic (GFAP, FABP7) markers in differentiated SREBF1 KO hiPSC-NSC clones and control cells (Cas9-only). Expression levels are normalized on the housekeeping gene GAPDH. Each dot represents the mean ± SEM of 2/3 technical replicates/clone. Mann-Whitney test: *, $p < 0.05$.

#### Supplementary Figure 7

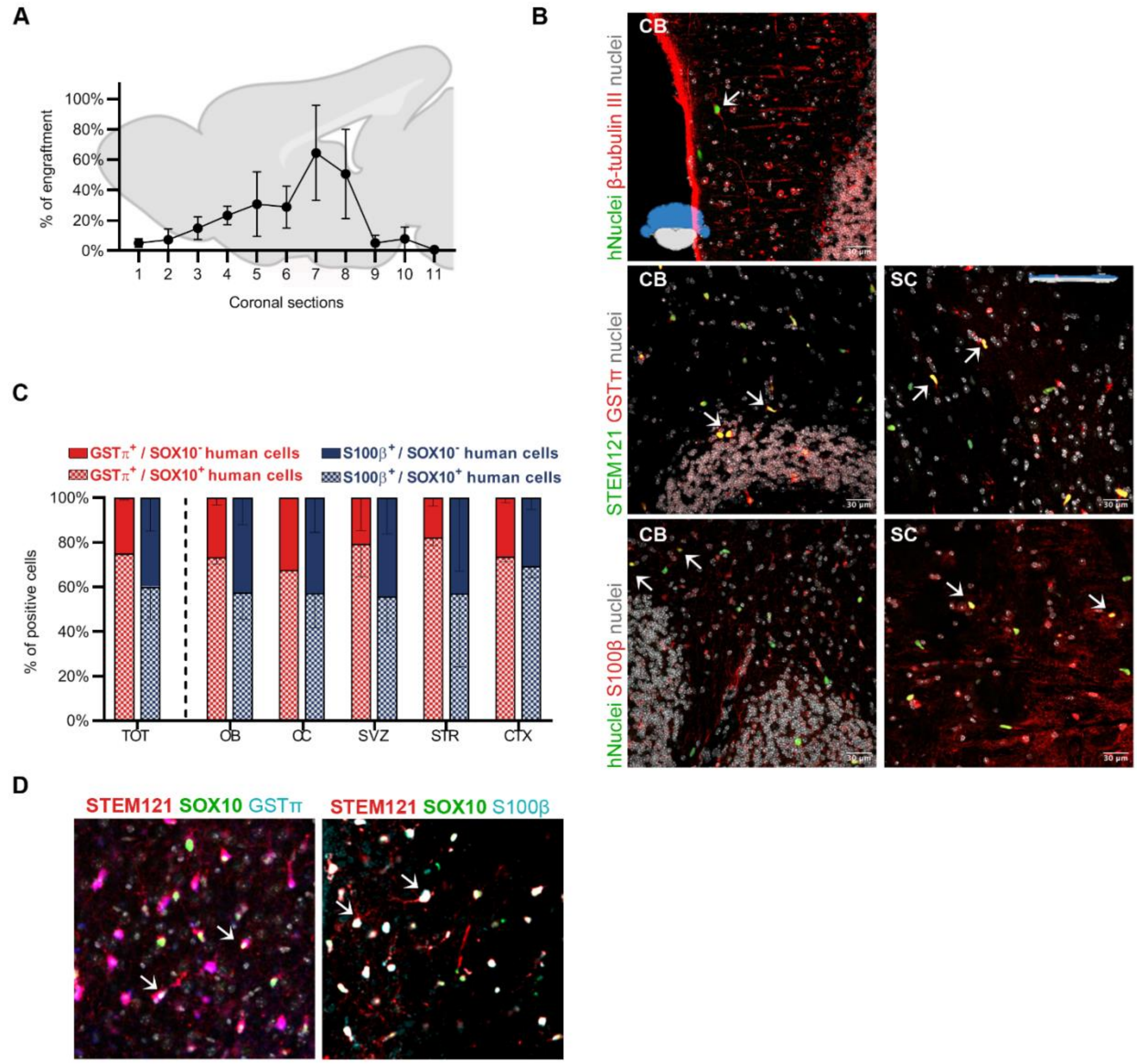


**Supplementary Figure 7. Engrafted hiPSC-NSCs migrate along the rostro-caudal axis and primarily differentiate into glia progenitors. A)** Graph showing the distribution of engrafted cells based along the rostro-caudal axis, evaluated as percentage of hNuclei$^+$ cells in sequential coronal sections. Data are represented as mean ± SEM (n = 2 mice). **B)** Representative immunofluorescence images of human cells (hNuclei$^+$ or STEM121$^+$) expressing S100β (astrocytes), GSTπ (oligodendrocytes), or β-tubulin III (neurons) markers in the cerebellum (CB) or spinal cord (SC). Nuclei were counterstained with Hoechst. Arrows indicate co-localization between immunofluorescence signals. **C)** Bar plot showing the percentage of hiPSC-NSC-derived GSTπ$^+$ (red bars) or S100β$^+$ (blue bars) cells co-expressing SOX10 marker (scattered bars). Percentages were calculated as: (number of STEM121$^+$GSTπ$^+$Sox10$^+$ or STEM121$^+$S100β$^+$Sox10$^+$ cells) / (number of STEM121$^+$GSTπ$^+$ or STEM121$^+$S100β$^+$ cells) × 100. Data are expressed as mean ± SEM (n = 6 mice). **D)** Representative immunofluorescence images of human cells (STEM121$^+$; red) co-expressing SOX10 (green) and GSTπ (blue) or SOX10 (green) and S100β (blue) (arrows).

#### Supplementary Tables

**Supplementary Table 1. Summary of hiPSC clones generated by reprogramming of healthy donor (HD) fibroblasts**. Skin fibroblasts derived from adult (HD1) and newborn (HD2) healthy donors were transduced with a monocistronic Cre-excisable lentiviral vector (LV) carrying OCT4, SOX2, and KLF4 under the control of the human SFFV promoter, as previously reported[34].

| ID | **Fibroblasts** (code number/source) | Age | hiPSC clones |
|---|---|---|---|
| HD1 | FFF0561980/ Gaslini Biobank | Adult | HD 1.1 |
| | | | HD 1.3 |
| HD2 | C0045C/Invitrogen | Newborn | HD 2.2 |
| | | | HD 2.3 |

**Supplementary Table 2. Upstream regulators identified by Ingenuity Pathway Analysis (IPA) on RNA-seq analysis of hiPSC-NPCs vs. hiPSCs**. IPA analysis identified the upstream regulators of genes up- or down-regulated during hiPSC neural commitment. Shown below are the gene name (Upstream Regulator), the inferred activation states of predicted transcriptional regulators (Predicted Activation State), activation *z*-score ($|z\text{-score}| > 2$) and the *p*-value of the overlap between the dataset genes and the genes that are regulated by an Upstream Regulator (*p*-value of overlap).

| Upstream Regulator | Predicted Activation State | Activation *z*-score | *p*-value of overlap |
|---|---|---|---|
| RBL1 | Activated | 2.556 | 9.06E-07 |
| FOXO3 | Activated | 2.514 | 6.49E-03 |
| MEF2D | Activated | 2.480 | 6.34E-04 |
| NEUROD1 | Activated | 3.261 | 5.92E-03 |
| CDKN2A | Activated | 3.688 | 5.54E-07 |
| KLF15 | Activated | 2.295 | 4.83E-02 |
| HEY2 | Activated | 2.006 | 4.56E-03 |
| SMARCB1 | Activated | 2.146 | 3.91E-08 |
| ZBTB17 | Activated | 2.219 | 3.90E-03 |
| HDAC1 | Activated | 3.425 | 3.25E-04 |
| KDM5B | Activated | 3.102 | 3.05E-06 |
| OTX2 | Activated | 2.550 | 2.73E-02 |
| TWIST1 | Activated | 2.018 | 2.72E-02 |
| MXD1 | Activated | 2.213 | 2.29E-03 |
| CREBBP | Activated | 2.339 | 2.24E-04 |
| EN1 | Activated | 2.213 | 2.23E-03 |
| ZEB2 | Activated | 2.080 | 2.06E-04 |
| PAX6 | Activated | 2.837 | 2.03E-02 |
| ZEB1 | Activated | 2.042 | 1.99E-03 |
| KDM5A | Activated | 2.194 | 1.90E-02 |
| NUPR1 | Activated | 3.580 | 1.71E-06 |
| HDAC2 | Activated | 3.115 | 1.64E-03 |
| SMARCA4 | Activated | 2.073 | 1.42E-06 |
| E2F6 | Activated | 2.688 | 1.25E-06 |
| FOXN4 | Activated | 2.236 | 1.21E-04 |
| TCF3 | Activated | 2.461 | 1.07E-02 |
| E2F1 | Inhibited | -4.041 | 1.89E-15 |
| MYCN | Inhibited | -3.976 | 2.60E-04 |
| POU5F1 | Inhibited | -3.589 | 9.00E-06 |
| CCND1 | Inhibited | -2.852 | 7.74E-15 |
| NANOG | Inhibited | -2.470 | 9.34E-06 |
| MYCBP | Inhibited | -2.000 | 1.69E-02 |
| MYC | Inhibited | -6.979 | 3.59E-10 |

| | | | |
|---|---|---|---|
| E2F3 | Inhibited | -4.397 | 7.50E-10 |
| TAL1 | Inhibited | -3.471 | 2.24E-05 |
| SMAD7 | Inhibited | -3.415 | 1.53E-02 |
| MAX | Inhibited | -2.902 | 1.83E-04 |
| TRIM24 | Inhibited | -2.828 | 2.37E-02 |
| REST | Inhibited | -2.698 | 2.30E-04 |
| BMI1 | Inhibited | -2.607 | 4.86E-04 |
| EPAS1 | Inhibited | -2.408 | 1.35E-05 |
| ZNF217 | Inhibited | -2.385 | 1.28E-06 |
| POU4F2 | Inhibited | -2.359 | 4.04E-05 |
| E2F2 | Inhibited | -2.325 | 2.89E-09 |
| MNT | Inhibited | -2.236 | 1.68E-03 |
| SIN3A | Inhibited | -2.236 | 3.58E-03 |
| ZIC2 | Inhibited | -2.219 | 8.50E-03 |
| HNF1B | Inhibited | -2.205 | 2.08E-04 |
| MED1 | Inhibited | -2.197 | 1.20E-07 |
| HAND2 | Inhibited | -2.139 | 2.36E-02 |
| TFAP2C | Inhibited | -2.094 | 6.53E-03 |
| HOXA10 | Inhibited | -2.063 | 3.41E-03 |
| MYCL | Inhibited | -2.000 | 3.19E-03 |

**Supplementary Table 3. Upstream regulators identified by Ingenuity Pathway Analysis (IPA) on the dataset of genes close to hiPSC- and hiPSC-NSC-specific enhancers**. IPA analysis of integrated ChIP-seq and RNA-seq datasets identified the upstream regulators of genes associated with gained and lost enhancers (400 kb window) in hiPSC-NSCs. Shown below are the gene name (Upstream Regulator), the inferred activation states of predicted transcriptional regulators (Predicted Activation State), activation *z*-score (│*z*-score│> 2) and the *p*-value of the overlap between the dataset genes and the genes that are regulated by an Upstream Regulator (*p*-value of overlap).

| **Upstream Regulator** | **Predicted Activation State** | **Activation *z*-score** | ***p*-value of overlap** |
|---|---|---|---|
| NONO | Activated | 2.711 | 4.17E-04 |
| SREBF1 | Activated | 3.215 | 6.41E-04 |
| STAT1 | Activated | 3.416 | 9.59E-04 |
| EHMT1 | Activated | 3.162 | 2.16E-02 |
| CTNNB1 | Activated | 2.425 | 2.38E-02 |
| RELA | Activated | 2.534 | 3.77E-02 |
| TEAD2 | Activated | 2.236 | 4.12E-02 |
| E2F1 | Inhibited | -3.022 | 9.23E-19 |
| ATF3 | Inhibited | -3.281 | 1.54E-10 |
| MYCN | Inhibited | -2.599 | 2.71E-09 |
| MYC | Inhibited | -4.925 | 2.37E-08 |
| E2F3 | Inhibited | -3.160 | 3.08E-08 |
| MITF | Inhibited | -4.914 | 8.61E-08 |
| TP63 | Inhibited | -3.261 | 2.87E-07 |
| YAP1 | Inhibited | -3.334 | 1.77E-06 |
| MYBL2 | Inhibited | -2.425 | 4.49E-05 |
| MED1 | Inhibited | -4.069 | 6.66E-05 |
| PAX8 | Inhibited | -2.392 | 8.77E-05 |
| TFAP2C | Inhibited | -2.178 | 1.23E-04 |
| HIF1A | Inhibited | -4.095 | 1.66E-04 |
| MYB | Inhibited | -2.615 | 2.94E-04 |

| | | | |
|---|---|---|---|
| SP1 | Inhibited | -2.559 | 3.51E-04 |
| FOXM1 | Inhibited | -3.917 | 4.40E-04 |
| NCOA3 | Inhibited | -3.136 | 4.61E-04 |
| EPAS1 | Inhibited | -2.931 | 6.68E-04 |
| TCF4 | Inhibited | -4.382 | 8.83E-04 |
| ZNF281 | Inhibited | -2.200 | 1.03E-03 |
| MYCBP | Inhibited | -2.000 | 1.20E-03 |
| FUS | Inhibited | -2.236 | 1.26E-03 |
| POU5F1 | Inhibited | -2.646 | 1.26E-03 |
| WWTR1 | Inhibited | -2.178 | 1.44E-03 |
| ID1 | Inhibited | -2.425 | 3.91E-03 |
| RELB | Inhibited | -2.376 | 4.58E-03 |
| GATA4 | Inhibited | -2.476 | 5.22E-03 |
| CREB1 | Inhibited | -2.813 | 7.60E-03 |
| KDM3A | Inhibited | -2.595 | 8.50E-03 |
| NFKB1 | Inhibited | -3.273 | 1.06E-02 |
| MYOCD | Inhibited | -2.466 | 1.31E-02 |
| NANOG | Inhibited | -2.359 | 1.38E-02 |
| EZH2 | Inhibited | -2.335 | 2.45E-02 |
| SRF | Inhibited | -2.219 | 2.79E-02 |
| CTNNB1 | Inhibited | -2.888 | 3.02E-02 |
| SMARCA4 | Inhibited | -4.526 | 3.30E-02 |
| POU2F2 | Inhibited | -3.162 | 3.43E-02 |
| STAT3 | Inhibited | -3.407 | 4.86E-02 |

**Supplementary Table 4. Upstream regulators identified by Ingenuity Pathway Analysis (IPA) on RNA-seq analysis of hiPSC-NSCs vs. hfNSCs**. IPA analysis identified the upstream regulators of genes upregulated in hiPSC-NSCs (*z*-score > 2) or hfNSCs (*z*-score < 2). Shown below are the gene name (Upstream Regulator), the inferred activation states of predicted transcriptional regulators (Predicted Activation State), activation *z*-score, and the *p*-value of the overlap between the dataset genes and the genes that are regulated by an Upstream Regulator (*p*-value of overlap).

| **Upstream Regulator** | **Predicted Activation State** | **Activation *z*-score** | ***p*-value of overlap** |
|---|---|---|---|
| NKX2-3 | Activated | 4.633 | 1.33E-11 |
| CTNNB1 | Activated | 4.249 | 3.83E-22 |
| NEUROG3 | Activated | 3.977 | 2.81E-04 |
| MYC | Activated | 3.853 | 1.14E-08 |
| TRIM24 | Activated | 3.429 | 2.66E-05 |
| KLF4 | Activated | 3.158 | 3.56E-08 |
| HIF1A | Activated | 3.123 | 1.41E-08 |
| SMAD3 | Activated | 3.098 | 3.52E-04 |
| HOXA9 | Activated | 3.065 | 3.12E-04 |
| SRF | Activated | 2.989 | 1.51E-05 |
| SOX11 | Activated | 2.784 | 4.04E-05 |
| ATF4 | Activated | 2.692 | 4.71E-04 |
| KMT2D | Activated | 2.571 | 1.79E-04 |
| HDAC6 | Activated | 2.547 | 1.07E-02 |
| NOTCH3 | Activated | 2.456 | 1.16E-02 |
| GFI1 | Activated | 2.397 | 5.45E-03 |
| HNF1A | Activated | 2.380 | 2.41E-03 |
| LEF1 | Activated | 2.355 | 2.50E-03 |
| GLI1 | Activated | 2.346 | 3.90E-09 |

| | | | |
|---|---|---|---|
| LHX1 | Activated | 2.340 | 1.17E-04 |
| STAT3 | Activated | 2.303 | 9.38E-09 |
| LMX1B | Activated | 2.219 | 2.85E-03 |
| SIM1 | Activated | 2.213 | 1.66E-06 |
| MAML1 | Activated | 2.173 | 2.53E-02 |
| SIX5 | Activated | 2.170 | 3.60E-04 |
| CDX1 | Activated | 2.164 | 4.44E-02 |
| CEBPB | Activated | 2.162 | 4.43E-02 |
| EGR2 | Activated | 2.158 | 5.51E-04 |
| SMAD4 | Activated | 2.101 | 1.65E-07 |
| ARNT2 | Activated | 2.101 | 1.35E-06 |
| FOXO1 | Activated | 2.030 | 1.81E-05 |
| OTX2 | Activated | 2.026 | 3.38E-04 |
| POU3F2 | Activated | 2.000 | 1.79E-02 |
| LMX1A | Activated | 2.000 | 1.27E-02 |
| GLIS1 | Activated | 2.000 | 2.35E-03 |
| IRF7 | Inhibited | -3.331 | 3.14E-04 |
| PAX1 | Inhibited | -3.207 | 6.77E-03 |
| TFEB | Inhibited | -3.153 | 4.73E-04 |
| NLRC5 | Inhibited | -3.113 | 2.77E-04 |
| REST | Inhibited | -3.010 | 5.73E-18 |
| STAT2 | Inhibited | -2.721 | 1.07E-03 |
| IRF1 | Inhibited | -2.658 | 1.46E-02 |
| PRDM8 | Inhibited | -2.646 | 7.30E-04 |
| SOX3 | Inhibited | -2.475 | 1.87E-09 |
| COMMD3-BMI1 | Inhibited | -2.379 | 1.74E-04 |
| TP53 | Inhibited | -2.254 | 2.19E-16 |
| HOXC9 | Inhibited | -2.236 | 1.79E-02 |
| GATA3 | Inhibited | -2.203 | 2.16E-02 |
| GMNN | Inhibited | -2.191 | 3.41E-07 |
| SOX1 | Inhibited | -2.191 | 2.04E-08 |
| SPDEF | Inhibited | -2.165 | 2.14E-07 |
| ZNF217 | Inhibited | -2.065 | 2.31E-07 |
| NFKB1 | Inhibited | -2.009 | 7.44E-03 |

**Supplementary Table 5. gRNA for SREBF1 KO and primers used for PCR amplification and sequencing.**

| **gRNA** | |
|---|---|
| gRNA1 | GAGCTCAAGGATCTGGTGGT |
| gRNA2 | TGCGCTTCTCTCCACGGCTC |
| gRNA3 | CGGAGAAGCTGCCTATCAAC |
| **Sequencing primers** | |
| FW | 5’-TAGCACAGCCCCACCTTTAT-3’ |
| Rev | 5’-AGCCATGAAGACAGACGGAG-3’ |

**Supplementary Table 6. List of primers used for qRT-PCR on immunoprecipitated chromatin.**

| **Primers for ChIP analysis** | |
|---|---|
| **Region** | **Sequence** |
| Negative | Fw 5’ -AAAGCTGGACTGGTGAATGC- 3’ |
| | Rev 5’ -TCAAAGGCTCATCTTTGCAG- 3’ |
| FGFR1 | Fw 5’ -GTCACAGCTGCCATCCTACA- 3’ |
| | Rev 5’ -TCTATTTGGGGACTCCGAGA- 3’ |
| Lin28 | Fw 5’ -CTCAGCAGTGGATGGGGATG- 3’ |

| | |
|---|---|
| | Rev 5’ -GCAGGAGGAACCCAAAGAGT- 3’ |
| PPAP2b | Fw 5’ -TGAGCATCGCTTTTCTGGGG- 3’ |
| | Rev 5’ -ACAGCTTGCTACGAGACAGG- 3’ |

**Supplementary Table 7. List of primers and probes used for SYBR Green and TaqMan qRT-PCR.**

| SYBR Green primers | |
|---|---|
| **Gene** | **Sequence** |
| PAX6 | Fw 5' -AGTGAATCAGCTCGGTGGTGTCTT- 3' |
| | Rev 5’ -TGCAGAATTCGGGAAATGTCGC- 3' |
| ROBO2 | Fw 5’ -TTCTTCTTGCGCATCGTGC- 3’ |
| | Rev 5’ -CGCATTTCGACTCACTGCTTC- 3’ |
| OCT4 | Fw 5' -TCGAGAACCGAGTGAGAGG- 3' |
| | Rev 5’ -GAACCACACTCGGACCACA- 3' |
| NANOG | Fw 5' -ATGCCTCACACGGAGACTGT- 3' |
| | Rev 5’ -AAGTGGGTTGTTTGCCTTTG- 3' |
| LIN28 | Fw 5’ -GAAGCGCAGATCAAAAGGAG- 3’ |
| | Rev 5’ -GCTGATGCTCTGGCAGAAGT- 3’ |
| **TaqMan probes** | |
| **Gene** | **Code** |
| PAX6 | Hs00240871_m1 |
| NEUROD1 | Hs01922995_s1 |
| MXD1 | Hs00965581_m1 |
| Max | Hs00231142_m1 |
| Myc | Hs00153408_m1 |
| SREBF1 | Hs01088691_m1 |
| ZEB1 | Hs00232783_m1 |
| GAPDH | Hs99999909_m1 |
| NKX2-2 | Hs00159616_m1 |
| NR3C2 | Hs01031804_m1 |
| GRHL1 | Hs01119372_m1 |
| MEIS1 | Hs00180020_m1 |
| PPARA | Hs00947536_m1 |
| CREB5 | Hs00191719_m1 |
| POU3F2 | Hs00271595_s1 |
| ZNF711 | Hs00944896_m1 |
| NPAS2 | Hs00231212_m1 |
| RXRB | Hs00232774_m1 |
| IRF2 | Hs01082884_m1 |
| JUN | Hs01103582_s1 |
| ARNT | Hs01121918_m1 |
| MEF2A | Hs01050406_g1 |
| FOXA1 | Hs04187555_m1 |
| NR3C1 | Hs00353740_m1 |
| TRIM3 | Hs01548703_m1 |
| ARNT2 | Hs00977663_m1 |
| HES1 | Hs00172878_m1 |
| NUPR1 | Hs01044304_g1 |
| FOXN4 | Hs01566111_m1 |
| FOXG1 | Hs01850784_s1 |
| DCX | Hs00167057_m1 |
| β-TUBULIN III | Hs00801390_s1 |
| MAP2 | Hs00258900_m1 |
| TH | Hs00165941_m1 |
| FABP7 | Hs00361424_g1 |
| GFAP | Hs00157674_m1 |

**Supplementary Table 8. List of primary and secondary antibodies with antigen, host species, provider, product number, and working dilutions indicated.**

| | **Primary Antibodies** | |
|---|---|---|
| **Antigen** | **Host species (provider, product number)** | **Working Dilution** |
| Human Nuclei | Mouse monoclonal (Sigma-Aldrich, MAB1281) | 1:100 |
| STEM121 | Mouse monoclonal (Takara Bio, Y40410) | 1:100 |
| GSTπ | Rabbit polyclonal (MBL, 312) | 1:500 |
| SOX10 | Goat polyclonal IgG (R&D Systems, AF2864) | 1:100 |
| β-tubulin III | Rabbit polyclonal IgG (BioLegend, 802001) | 1:2000 |
| S100 β | Rabbit (Swant, 37A) | 1:1000 |
| Human Nestin | Rabbit polyclonal (Millipore, ABD69) | 1:200 |
| Ki67 (D3B5) | Rabbit monoclonal IgG (Cell Signaling, MAB9129) | 1:200 |
| Ki67 | Mouse polyclonal (Novocastra, NCL-Ki67-MM1) | 1:100 |
| | **Secondary antibodies** | |
| Alexa 488 | Goat anti-Mouse IgG (Mol Probes, A11001) | 1:1000 |
| Alexa 546 | Goat anti-Rabbit IgG (Mol Probes, A11010) | 1:2000 |
| | **Nuclear Counterstain** | |
| Hoechst 33342 | (Invitrogen, H3570) | 1:1000 |
